# Dual Impacts of a Glycan Shield on the Envelope Glycoprotein B of HSV-1: Evasion from Human Antibodies In Vivo and Neurovirulence

**DOI:** 10.1101/2022.12.12.519993

**Authors:** Ayano Fukui, Yuhei Maruzuru, Shiho Ohno, Moeka Nobe, Shuji Iwata, Kosuke Takeshima, Naoto Koyanagi, Akihisa Kato, Shinobu Kitazume, Yoshiki Yamaguchi, Yasushi Kawaguchi

**Affiliations:** Division of Molecular Virology, Department of Microbiology and Immunology, The Institute of Medical Science, The University of Tokyo, Tokyo, Japan; Department of Infectious Disease Control, International Research Center for Infectious Diseases, The Institute of Medical Science, The University of Tokyo, Tokyo, Japan; Research Center for Asian Infectious Diseases, The Institute of Medical Science, The University of Tokyo, Tokyo, Japan; Division of Structural Glycobiology, Institute of Molecular Biomembrane and Glycobiology, Tohoku Medical and Pharmaceutical University, Miyagi, Japan; Department of Clinical Laboratory Sciences, School of Health Sciences, Fukushima Medical University, Fukushima, Japan

## Abstract

Identification of the mechanisms of viral evasion from human antibodies is crucial both for understanding viral pathogenesis and for designing effective vaccines. However, the in vivo efficacy of the mechanisms of viral evasion from human antibodies has not been well documented. Here we show in cell cultures that an N-glycan shield on the HSV-1 envelope glycoprotein B (gB) mediated evasion from neutralization and antibody-dependent cellular cytotoxicity due to pooled γ-globulins derived from human blood. We also demonstrated that the presence of human γ-globulins in mice and HSV-1 immunity induced by viral infection in mice significantly reduced the replication of a mutant virus lacking the glycosylation site in a peripheral organ but had little effect on the replication of its repaired virus. These results suggest that the glycan shield on the HSV-1 envelope gB mediated evasion from human antibodies in vivo and from HSV-1 immunity induced by viral infection in vivo. Notably, we also found that the glycan shield on HSV-1 gB was significant for HSV-1 neurovirulence and replication in the central nervous system (CNS) of naïve mice. Thus, we have identified a critical glycan shield on HSV-1 gB that has dual impacts, namely evasion from human antibodies in vivo and viral neurovirulence.

**IMPORTANCE:** HSV-1 establishes lifelong latent and recurrent infections in humans. To produce recurrent infections that contribute to transmission of the virus to new human host(s), the virus must be able to evade the antibodies persisting in latently infected individuals. Here we show that an N-glycan shield on the envelope glycoprotein B of HSV-1 mediates evasion from pooled γ-globulins derived from human blood both in cell cultures and mice. Notably, the N-glycan shield was also significant for HSV-1 neurovirulence in naïve mice. Considering the clinical features of HSV-1 infection, these results suggest that the glycan shield not only facilitates recurrent HSV-1 infections in latently infected humans by evading antibodies, but is also important for HSV-1 pathogenesis during the initial infection.

## INTRODUCTION

Herpes simplex viruses (HSV)-1 and HSV-2 cause a variety of human diseases, including encephalitis; keratitis; neonatal disease; and mucocutaneous and skin diseases such as herpes labialis, genital herpes, and herpetic whitlow (1–3). A striking feature of these viruses is that they establish lifelong infections in humans, where, after the initial infection, they become latent and frequently reactivate to cause lesions (1–3). To accomplish these cycles, the viruses have evolved highly complex and sophisticated strategies to evade host immune mechanisms. Notably, these viral strategies have probably impeded the development of effective vaccines for HSV-1 and HSV-2 infections. Several decades of vaccine development have not produced a successful vaccine (4, 5).

To clarify the significance of the mechanisms of immune evasion by viruses that cause diseases in humans, the mechanisms should be investigated not only in vitro but also in vivo; and research using available human samples should provide valuable information on the effective viral mechanisms in humans. However, in previous studies, human samples have generally been analyzed in vitro, and information from the in vivo evaluations of human samples on viral evasion from the immune system has been limited. To fill the gaps in our understanding of the effective mechanisms of viral immune evasion in humans, in vivo investigations of the mechanisms of immune evasion that use human samples are of crucial importance.

Although no effective vaccines for HSV-1 and HSV-2 have been developed thus far, previous clinical trials for HSV vaccines have provided important clues indicating that not only T-cell responses, but also antibody responses were important for controlling HSV infections in humans (4, 5). Thus, a clinical trial of a subunit vaccine employing HSV-2 envelope glycoprotein D (gD) showed 82% efficacy against the development of HSV-1 genital disease but did not offer significant protection against HSV-2 genital disease (6). Notably, antibody responses to HSV-2 gD correlated with protection against HSV-1 but not HSV-2 infections, whereas CD4^+^ T-cell responses did not correlate with protection against either HSV-1 or HSV-2 infection (7). In addition, a substudy of this trial that used sera from a fraction of the vaccinated subjects showed that neutralizing antibody titers against HSV-1 were significantly higher than the titers against HSV-2 (8). These findings were in agreement with those from another clinical study in humans that showed that the absence of HSV antibodies was associated with severe HSV infections in humans (9).

The findings described in the previous paragraph suggesting that antibodies are important for the control of HSV-1 infections led us to attempt to identify hitherto unknown mechanisms of HSV-1 evasion from human antibodies. In this study, we focused on the glycosylation of a major envelope glycoprotein of HSV-1, gB. Glycosylation of a viral envelope glycoprotein sometimes acts as a glycan shield for evading antibodies (10). The HSV-1 gB is a major target of antibody-mediated immunity (11). HSV-1 gB, which is a class III fusion glycoprotein, plays an essential role in the entry of the virus into a host cell, together with other HSV-1 envelope glycoproteins, including gD and a complex of gH and gL (gH/gL) (12). Herein we investigated effects of a series of N-linked glycans (N-glycans) on HSV-1 gB in the context of viral infection, and identified an N-glycan that contributed to evasion from human antibodies not only in vitro but also in vivo. Notably, the N-glycan on gB was also significant for HSV-1 replication in the central nervous system (CNS) of naïve mice as well as neurovirulence, although it had no effect on viral replication in cell cultures.

## RESULTS

### Generation of recombinant viruses harboring a mutation in each of the potential N-glycosylation sites on HSV-1 gB by an improved genetic manipulation system for HSV-1

HSV-1 gB has 6 potential N-glycosylation sites at the following positions: Asn-87, -141, -398, -430, -489, and -674 (Fig. 1). To investigate the significance of N-glycosylation on HSV-1 gB in the context of viral infection, we used an improved HSV-1 genetic manipulation system to construct a series of recombinant viruses and their repaired viruses (S-Fig. 1). The recombinant viruses encoded mutant gBs (gB-N87Q, - N141Q, -N398Q, -N430Q, -N489Q, and -N674Q), in which each of the potential N-glycosylation sites was substituted with glutamine. In addition, we generated a pair of control viruses, a recombinant virus, in which Asn-888 in the cytoplasmic domain of gB was substituted with glutamine (gB-N888Q), and its repaired virus (gB-N888Q-repair) (S-Fig. 1).

**Fig. 1.**
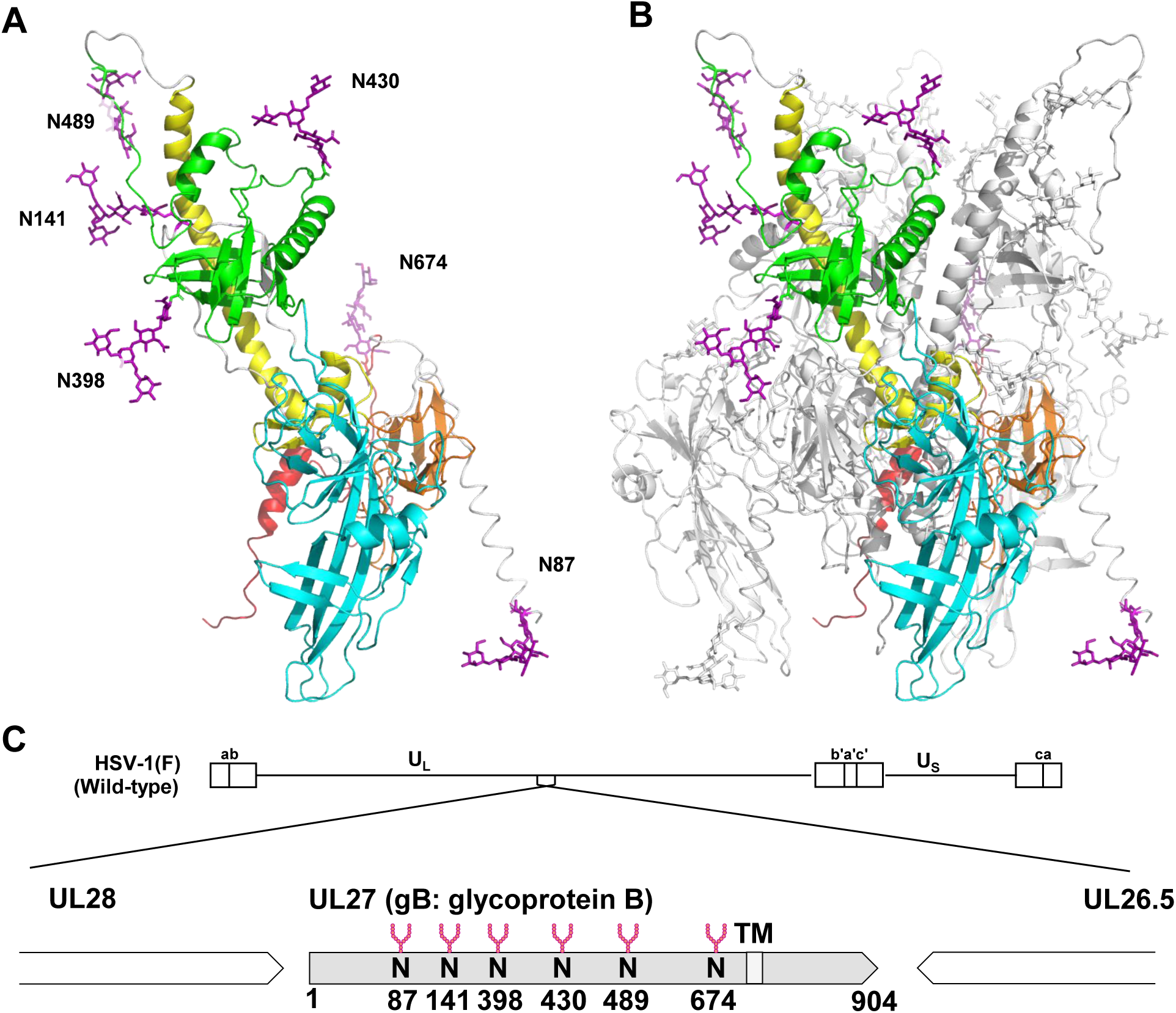
A 3D structural model of fully N-glycosylated gB in the prefusion state. Image A shows the ribbon diagram of the protomer, and image B shows the trimer of the crystal structure of gB in the prefusion state (Protein Data Bank [PDB] accession no. 6Z9M) (18). The Glycan Reader and Modeler were used to modify potential N-glycosylation sites (Asn-87, Asn-141, Asn-398, Asn-430, Asn-489, and Asn-674) of prefusion gB with the Man_3_GlcNAc_2_ glycan using Glycan Modeler. Missing coordinates of Asn-87 and Asn-489 in the 3D structure were estimated. The ribbon diagrams of the gB models show the functional domains in colors as follows: domain I (light blue), domain II (green), domain III (yellow), domain IV (orange), and domain V (red). Residue background coloring is used for the main polypeptide chains, and potential N-glycans are shown in magenta as stick models. In Panel B, protomer A is the same as in Panel A. Protomer B and C are shown in white. Image C shows the location of potential N-linked glycans on gB along the genome of wild-type HSV-1(F). N, N-glycosylation sites (Asn-87, Asn-141, Asn-398, Asn-430, Asn-489 and Asn-674); TM, transmembrane domain.

The two-step Red-mediated recombination system consists of the first recombination for the insertion of a PCR-amplified selectable marker and the second recombination for the excision of the inserted marker by a cleavage step that uses a rare-cutting endonuclease I-SceI. This system is widely used for markerless modifications of large DNA molecules such as herpesvirus genomes that are cloned into a bacterial artificial chromosome (BAC) in *Escherichia coli* (13).

However, the second recombination step was not very efficient in our hands. Therefore replica-plating, which is time-consuming and laborious, was needed to identify any *E. coli* harboring a BAC clone with a desired mutation. We used an improved system that was developed for this study that employed a negative selection marker, the *E. coli* phenylalanyl-tRNA synthetase (ePheS*), which encodes a mutant of the α-subunit of *E. coli* phenylalanyl-tRNA synthetase (ePheS) (14) in the presence of 4-chloro-phenylalanine (4CP), in addition to cleavage by I-SceI for the second recombination step (S-Fig. 2A). The improved system considerably increased the efficiency of the second recombination step from 17.4% to 87%, and 17.4 to 94.7% and 8.7% to 87.0% in the substitution of single amino acids, and in the deletion and insertion of a foreign gene, respectively, compared with the original system (S-Table 1A). Recombinant viruses carrying an alanine substitution of Thr-190 in the HSV-1 protein UL51 (S-Fig. 3A) that were generated in the original and improved systems exhibited identical growth properties (S-Fig. 3B) in Vero cells and produced identical neurovirulence in mice following intracranial inoculation (S-Fig. 3C), suggesting that, compared with the original system, the additional negative selection in the improved system did not affect the genomic integrity of HSV-1 other than the desired mutations.

### Effects of mutations at each of the potential gB N-glycosylation sites on electrophoretic mobility in the presence or absence of peptide-N-glycosidase F (PNGase)

Vero cells infected with wild-type HSV-1(F), each of the gB mutant viruses, or each of their repaired viruses were lysed, treated with or without PNGase, and analyzed by immunoblotting. As shown in Fig. 2, all of the gB mutants, except the gB-N888Q mutant, migrated faster than wild-type gB in denaturing gels. In contrast, the gB-N888Q mutant and gB from cells infected with each of the repaired viruses migrated as slowly as the wild-type gB in denaturing gels (Fig. 2). After treatment of the infected cell lysates with PNGase, all of the gB mutants migrated in denaturing gels as slowly as the wild-type gB (Fig. 2). All of the gB mutants were detected by immunoblotting at levels similar to the level of wild-type gB (Fig. 2). These results indicate that gB was N-glycosylated at each of the 6 potential N-linked glycosylation sites without affecting its accumulation in HSV-1-infected cells.

**Fig. 2.**
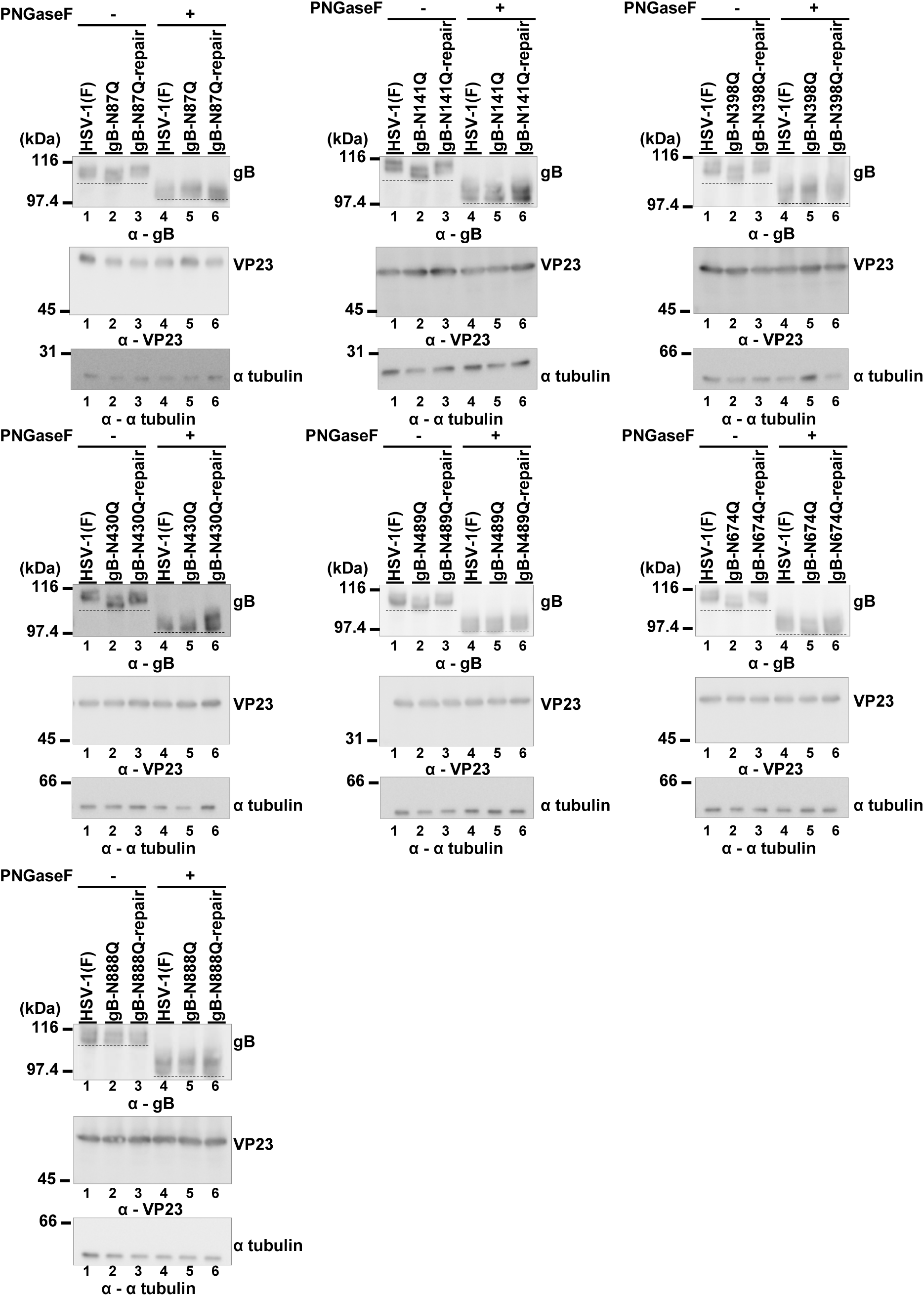
Effect of mutation at each of the potential gB N-glycosylation sites on electrophoretic mobility in the presence or absence of PNGase. Vero cells infected for 24 h with wild-type HSV-1(F), each of the gB mutant viruses, or each of their repaired viruses at an MOI of 5 were lysed, treated with or without PNGase, and analyzed by immunoblotting with antibodies to gB, VP23, or α-tubulin. Data are representative of 3 independent experiments. Dashed lines indicate the bottom of bands harboring gB with each of the indicated NQ mutations. α, anti.

### Effects of mutations at each of the gB N-glycosylation sites on the replication of HSV-1 in cell cultures

To investigate the effect of gB N-glycosylation on HSV-1 replication in cell cultures, Vero cells were infected with wild-type HSV-1(F), each of the gB mutant viruses, or each of their repaired viruses at a multiplicity of infection (MOI) of 5 or 0.01; and virus titers were assayed at 24 or 48 h postinfection. As shown in S-Fig. 4, progeny virus yields in cells infected with each of the gB mutant viruses were similar to those in cells infected with wild-type HSV-1(F) or each of their repaired viruses. These results suggest that N-glycosylation on gB did not affect HSV-1 replication in cell cultures.

### Effects of mutations at each of the gB N-glycosylation sites on viral susceptibility to neutralization by human antibodies

An estimated 67% of the global human population is infected with HSV-1 (15). Therefore, we decided to use pooled γ-globulins from human blood that contain amounts of antibodies to HSV-1 sufficient for our experiments (16). Indeed, we showed that gB antibodies in pooled human γ-globulins at a concentration of 1.3 mg/mL could still be detected at a dilution of 1:1,024 (S-Fig. 5). To investigate the effect of gB N-glycosylation on viral susceptibility to neutralization by human antibodies, the sensitivity to neutralization of wild-type HSV-1(F) by pooled human γ-globulins was compared to each of the recombinant gB mutant viruses. Among the gB mutant viruses tested, gB-N141Q was only the gB mutant virus that was significantly more susceptible to neutralization by pooled human γ-globulins at a concentration of 0.041 mg/mL compared with wild-type HSV-1(F) (S-Fig. 6). Therefore, we focused on N-glycosylation at gB Asn-141 and further characterized gB-N141Q in detail.

At pooled human γ-globulin concentrations ranging from 0.010 to 0.041 mg/mL, gB-N141Q was significantly more susceptible to neutralization by the human γ-globulins than wild-type HSV-1(F) (Fig. 3A). Wild-type susceptibility was restored in the gB-N141Q-repair virus (Fig. 3A). Antibodies to gB or gD in the human γ-globulins (0.082 mg/ml) were then depleted by treatment of the human γ-globulins with purified gB or gD fused with Strep-tag at the C-terminus (gB-SE or gD-SE, respectively) (S-Fig. 7A and B). The anti-gB antibody-depleted human γ-globulins could not detect gB ectopically expressed by HEK293FT cells (S-Fig. 7C and D). As shown in Fig. 3B, the susceptibility of gB-N141Q to neutralization by anti-gB antibody-depleted human γ-globulins (∼0.4 mg/mL) was comparable to that of wild-type HSV-1(F) and gB-N141Q-repair. In contrast, gB-N141Q was significantly more susceptible to neutralization by mock-depleted or anti-gD antibody-depleted human γ-globulins than wild-type HSV-1(F) and gB-N141Q-repair (Fig. 3B) as observed in Fig. 3A, showing its susceptibility to human γ-globulins without depletion. These results suggest that the N-glycan on gB Asn-141 was required for efficient HSV-1 evasion from neutralization by human antibodies that targeted gB in cell cultures.

**Fig. 3.**
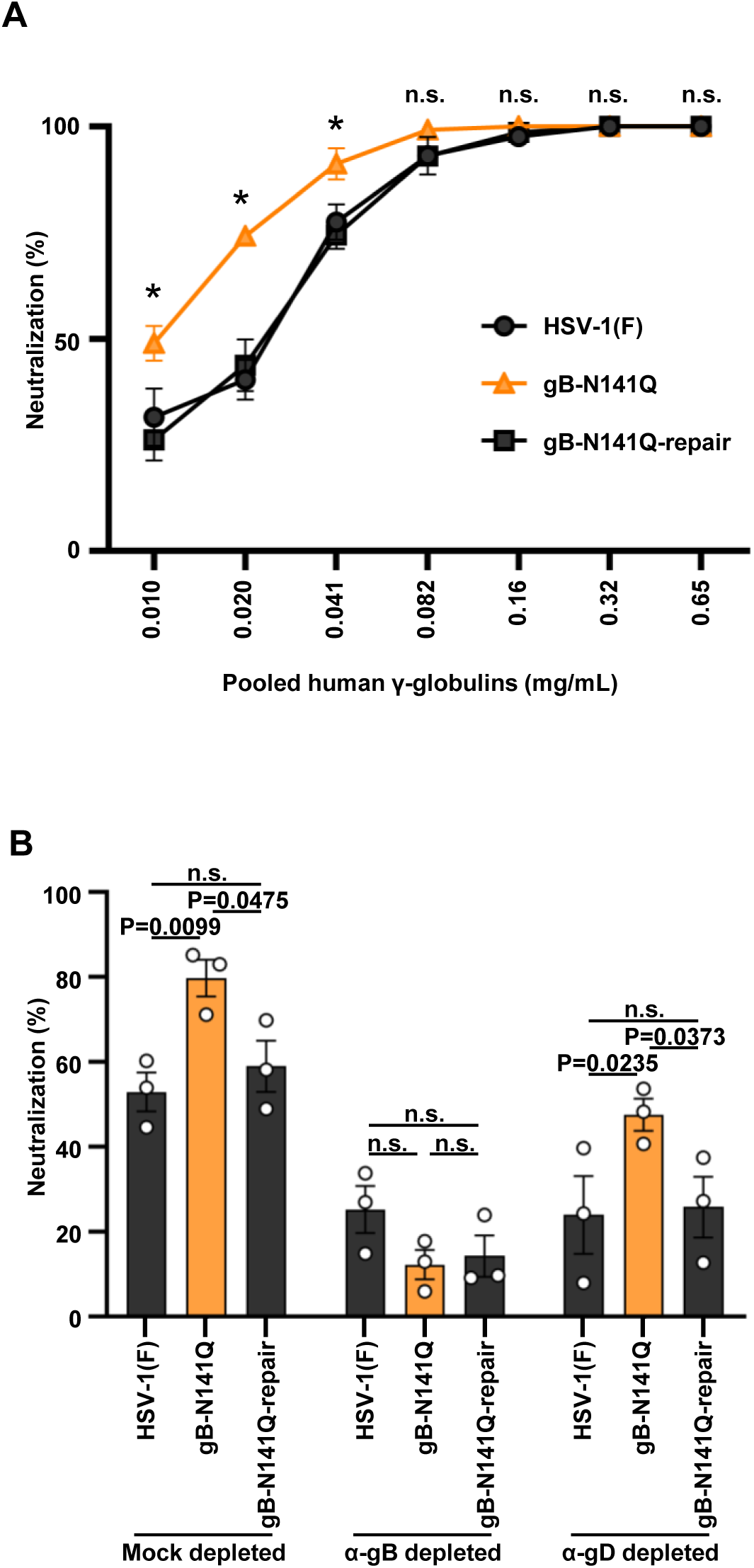
Effect of N-glycan at gB Asn-141 on viral susceptibility to neutralization by pooled human γ-globulins. (A) 100 PFU of wild-type HSV-1(F), gB-N141Q, or gB-N141Q-repair were incubated with serially diluted human γ-globulins at 37°C for 1 h, and then inoculated onto Vero cell monolayers for plaque assays. The percentage of neutralization was calculated from the number of plaques formed by each of the viruses that were incubated with or without human γ-globulins, as follows: 100×[1-(number of plaques after incubation with human γ-globulins)/(number of plaques after incubation without human γ-globulins)]. (B) Human γ-globulins (0.082 mg/mL) were mock-depleted (mock-depleted) or depleted with gB-SE (α-gB depleted) or gD-SE (α-gD depleted) as shown in S-Fig. 7. Wild-type HSV-1(F), gB-N141Q, or gB-N141Q-repair were incubated with each of the depleted human γ-globulins and then inoculated onto Vero cell monolayers as described in A. Each value represents the mean ± standard error of the results of 3 independent experiments. Statistical analysis was performed by two-way ANOVA followed by the Tukey test (A and B). *, *P* < 0.05 indicates statistically significant differences between gB-N141Q and wild-type HSV-1(F) or gB N141Q-repair; n.s., not significant; α, anti.

### Effects of the N-glycan on gB Asn-141 on human antibody-dependent cellular cytotoxicity (ADCC)

It has been reported that gB on the surface of infected cells mediates ADCC (17); therefore, we examined the effect of the N-glycan on gB Asn-141 on ADCC induced by human γ-globulins. Vero cells infected with wild-type HSV-1(F), gB-N141Q, or gB-N141Q-repair at an MOI of 1 for 24 h were subjected to an activating FcγRIIIA receptor ADCC assay in the presence or absence of human γ-globulins. As shown in Fig. 4A, human γ-globulins at concentrations 0.33 and 1.0 1mg/mL induced significantly higher FcγRIIIA activation in cells infected with gB-N141Q than in cells infected with wild-type HSV-1(F) or gB-N141Q-repair. The gB and gD antibodies present in samples of human γ-globulins at a concentration of 3.04 mg/mL were then depleted by treatment with purified gB-SE or gD-SE, as described previously (S-Fig. 7). In this case, the anti-gB antibody-depleted human γ-globulins slightly detected the gB that was ectopically expressed by HEK293FT cells (S-Fig. 7E), because we used the depleted human γ-globulins at a much higher concentration than the concentration of the depleted human γ-globulins used in the neutralizing assay described in the previous section. Consistent with the results shown in S-Fig. 7E, anti-gB antibody-depleted human γ-globulins (∼1.0 mg/mL) still induced slightly increased FcγRIIIA-mediated activation in cells infected with gB-N141Q than in cells infected with wild-type HSV-1(F) or gB-N141Q-repair (Fig. 4B). However, the degrees of differences between the levels of FcγRIIIA-mediated activation in cells infected with gB-N141Q and the levels of FcγRIIIA-mediated activation in cells infected with wild-type HSV-1(F) or gB-N141Q-repair in cultures containing anti-gB antibody-depleted human γ-globulins were lower than the degrees of differences between the levels of FcγRIIIA-mediated activation in cells infected with those viruses in cultures containing mock-depleted or anti-gD antibody-depleted human γ-globulins (Fig. 4B).

**Fig. 4.**
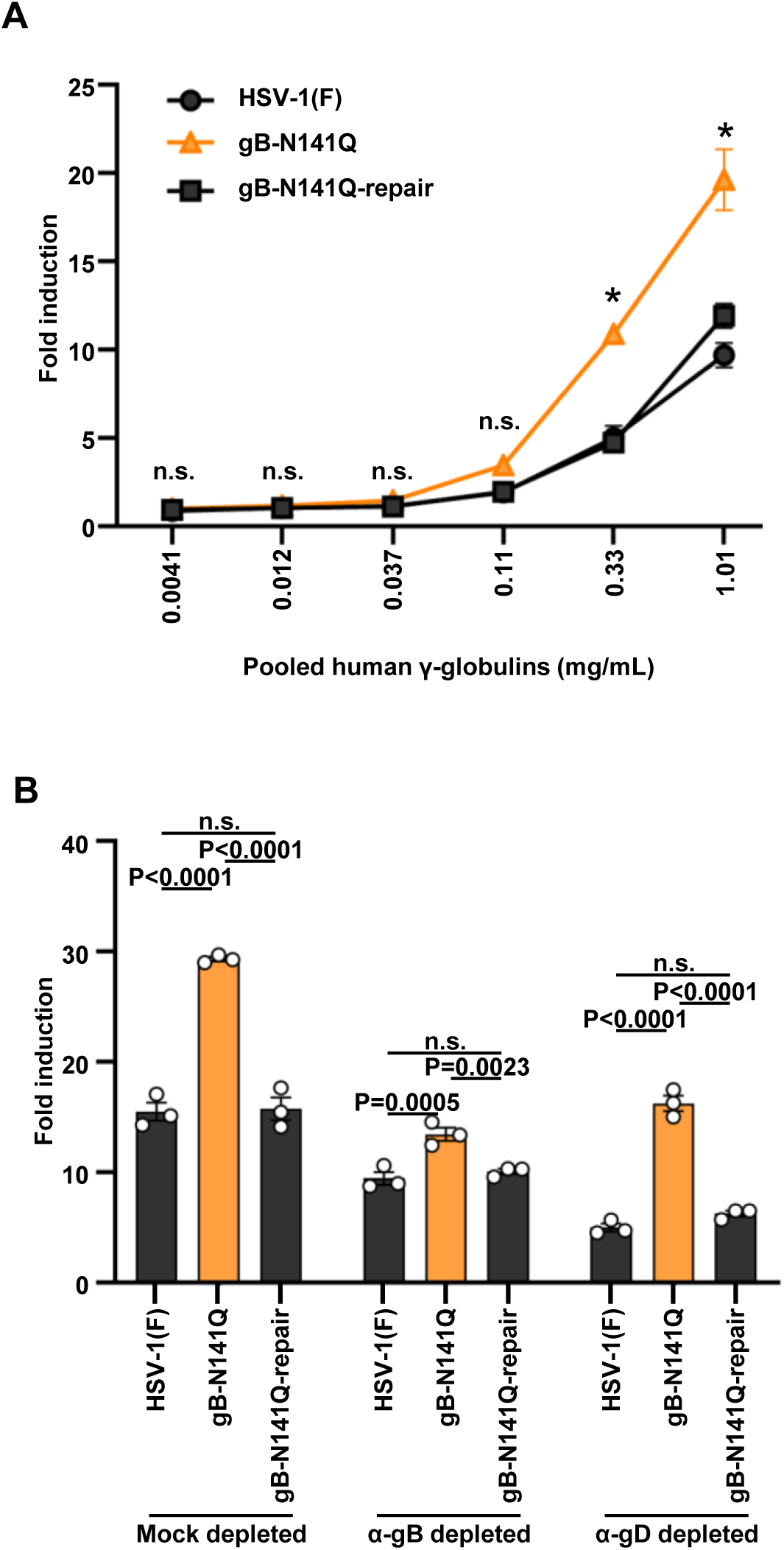
Effects of N-glycan at Asn-141 in gB on the extent of ADCC induced by human γ-globulins. (A) Vero cells were infected with wild-type HSV-1(F), gB-N141Q or gB-N141Q-repair at an MOI of 1 for 24 h, and co-cultured with ADCC effector cells in the presence or absence of serially diluted human γ-globulins for 6 h. A luciferase assay was then performed. (B) Human γ-globulins (3.04 mg/mL) were mock-depleted (mock-depleted) or depleted with gB-SE (α-gB depleted) or gD-SE (α-gD depleted) as depicted in S-Fig. 7 and used as described in 4A. Values are fold induction relative to controls without antibody. Each value is the mean ± standard error of the results of 3 biologically independent samples. The statistical analysis was performed by two-way ANOVA followed by the Tukey test. *, *P* < 0.05 indicates statistically significant differences between gB-N141Q and wild-type HSV-1(F) or gB-N141Q-repair; n.s., not significant; α, anti.

To eliminate the possibility that the higher level of FcγRIIIA-mediated activation in gB-N141Q-infected cells was due to increased expression of mutated gB on the surface of the infected cells, we investigated the effect of the N-glycan at gB Asn-141 on the expression of gB in the infected cells. Vero cells were infected with wild-type HSV-1(F), gB-N141Q, gB-N141Q-repair, ΔgB, or ΔgB-repair as described in the experiments described in the previous paragraph and depicted by Fig. 4 and used flow cytometry to show the level of gB expression on the surface of infected cells or the total accumulation of gB in the infected cells. As shown in S-Figs. 8A and B, the levels of expression of gB on the surface of cells infected with gB-N141Q were significantly lower than those levels on cells infected with wild-type HSV-1(F) or gB-N141Q-repair. In contrast, the total level of gB in cells infected with gB-N141Q was similar to the total levels in cells infected with wild-type HSV-1(F) or gB-N141Q-repair (S-Fig. 8A). Furthermore, confocal microscopy showed that the subcellular localization of gB in cells infected with either gB-N141Q, wild-type HSV-1(F) or gB-N141Q-repair was also similar (S-Fig. 8C). These results suggest that the N-glycan at gB Asn-141 was required for the efficient expression of gB on the surface of HSV-1-infected cells. Thus, although cells infected with gB-N141Q expressed lower levels of mutated gB on their surface membranes than the levels of gB expressed by cells infected with wild-type HSV-1(F) or gB-N141Q-repair, human γ-globulins resulted in increased FcγRIIIA-mediated activation of cells infected with gB-N141Q than seen for cells infected with wild-type HSV-1(F) or gB-N141Q-repair. These results eliminated the possibility that the higher level of FcγRIIIA-mediated activation in gB-141Q-infected cells was due to the increased expression of mutated gB on the surface of the infected cells. Altogether, the results suggest that the N-glycan at gB Asn-141 was required for the efficient evasion of ADCC induced by human antibodies to gB in infected cell cultures.

### Molecular modeling of N-glycosylated gB to estimate the glycan shield

To estimate the effects of the N-glycan on gB Asn-141 on the binding of antibodies to gB, we first constructed a glycosylated model of the gB protein based on a previously published construction of gB in a prefusion state (18). The Man_3_GlcNAc_2_ glycan structure was chosen for modeling. It is the common core structure of complex, high-mannose- and hybrid-type N-glycans.

The extent of antibody accessibility to gB was then estimated by determining the accessible surface area (ASA) with the use of a probe with a radius of 10 Å, which should be sufficient for determining the area of an antibody complementarity-determining region (19). The effect of N-glycosylation on gB Asn-141 on the ASA of each amino acid residue was evaluated by the difference between the ASAs (ΔASA) of N-glycosylated and non-glycosylated gB. Twenty-seven amino acid residues in the gB of HSV-1 showed ΔASAs of > 5 Å^2^ (S-Fig. 9) and as depicted by the 3D structural model of N-glycosylated gB in Fig. 5A. Among the 27 amino acid residues, 19 were mapped to the functional region (FR)2 and FR3 of gB (S-Fig. 9), both of which were previously defined based on the epitopes seen for a panel of neutralizing monoclonal antibodies (20, 21). Of note, the ΔASA of Asp-419, part of the epitope for the C226 antibody and critical for binding of that antibody to gB (21), measured 14, 40, and 52 Å^2^, depending on each protomer of the gB triplex (Fig. 5B and S-Fig. 9). These results suggest that the N-glycan at gB Asn-141 prevented the binding of antibodies to gB epitopes and further supported our previous conclusion that the N-glycan at gB Asn-141 was required for the efficient evasion of HSV-1 from human antibodies.

**Fig. 5.**
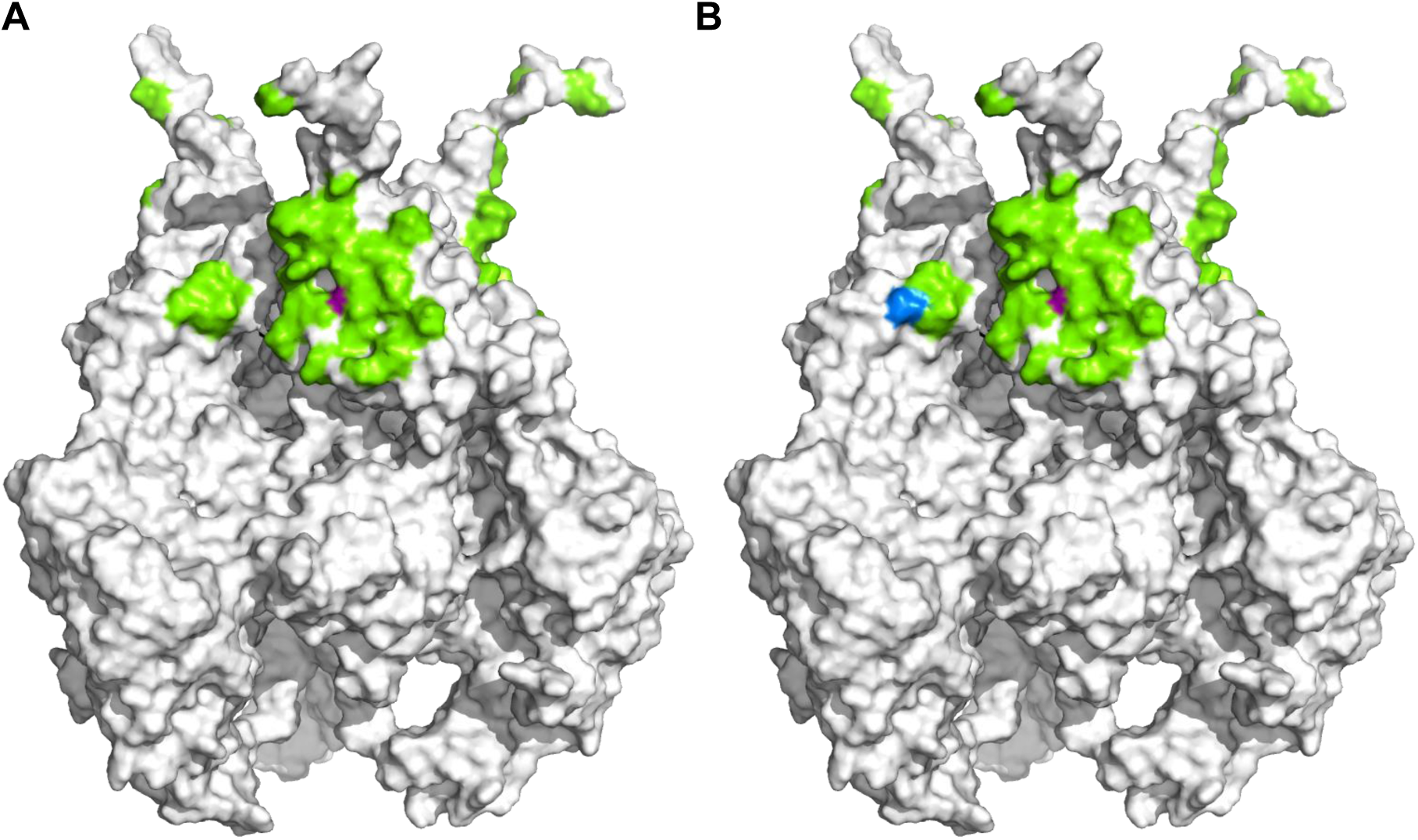
Effects of N-glycan shield at Asn-141 in gB on the antigenicity of gB in the prefusion state. (A) Mapping of amino acid residues potentially shielded by the N-glycan at Asn-141 in gB. The amino acid residues with ΔASAs greater than 5 Å^2^ are colored in green on the 3D structural model of the gB trimer in the prefusion state. Asn-141 is shown in magenta. (B) Overlapping of the C226 epitope which is targeted by neutralizing antibodies to gB and the glycan-shielded region. The amino acid residue with an ΔASA greater than 5 Å^2^ that also overlaps with the epitope of the neutralizing antibody C226 is shown in blue. The amino acid residues showing ΔASAs greater than 5 Å^2^ that do not overlap with the amino acid residues in the epitopes of the antibody are shown in green. Asn-141 is shown in magenta.

### Effects of the N-glycan on gB Asn-141 on the replication of HSV-1 in the eyes of mice in the presence of human antibodies

To examine the effects of human antibodies on HSV-1 replication in vivo in the presence or absence of the N-glycan at gB Asn-141, mice were mock-injected or injected intraperitoneally with pooled human γ-globulins, and were then ocularly infected with gB-N141Q or gB-N141Q-repair one day after injection (Fig. 6A). Samples of tear films were collected at the indicated times (Fig. 6A) and viral titers in the tear films were measured. As shown in Fig. 6B, the presence of human γ-globulins did not affect the viral titers of the tear films in mice infected with gB-N141Q-repair at 1, 3, and 5 days postinfection. In contrast, the presence of human γ-globulins significantly reduced viral titers of the tear films of mice infected with gB-N141Q at 1 and 5 days postinfection (Fig. 6B). Thus, the ratios of gB-N141Q titers in the absence of human γ-globulins to those in the presence of human γ-globulins were higher than the ratios of gB-N141Q-repair titers in the absence of human γ-globulins to those in the presence of the human γ-globulins (Fig. 6C). Furthermore, viral titers of the tear films of mice infected with gB-N141Q in the presence of human γ-globulins at 1, 3, and 5 days postinfection were significantly lower than the titers of the tear films of mice infected with gB-N141Q-repair (Fig. 6B). In contrast, the viral titers of the tear films of mice infected with gB-N141Q in the absence of human γ-globulins at 1, 3, and 5 days postinfection were comparable to those of the tear films of mice infected with gB-N141Q-repair; although, as the infection progressed, the viral titers of the tear films of mice infected with gB-N141Q in the absence of human γ-globulins tended to be lower than those titers in mice infected with gB-N141Q-repair. These results indicate that the presence of human antibodies inhibited the replication of gB-N141Q in the peripheral organs of mice more efficiently than it inhibited the replication of gB-N141Q-repair and also suggest that the N-glycan at gB Asn-141 was required for the efficient evasion of HSV-1 from human antibodies in vivo.

**Fig. 6.**
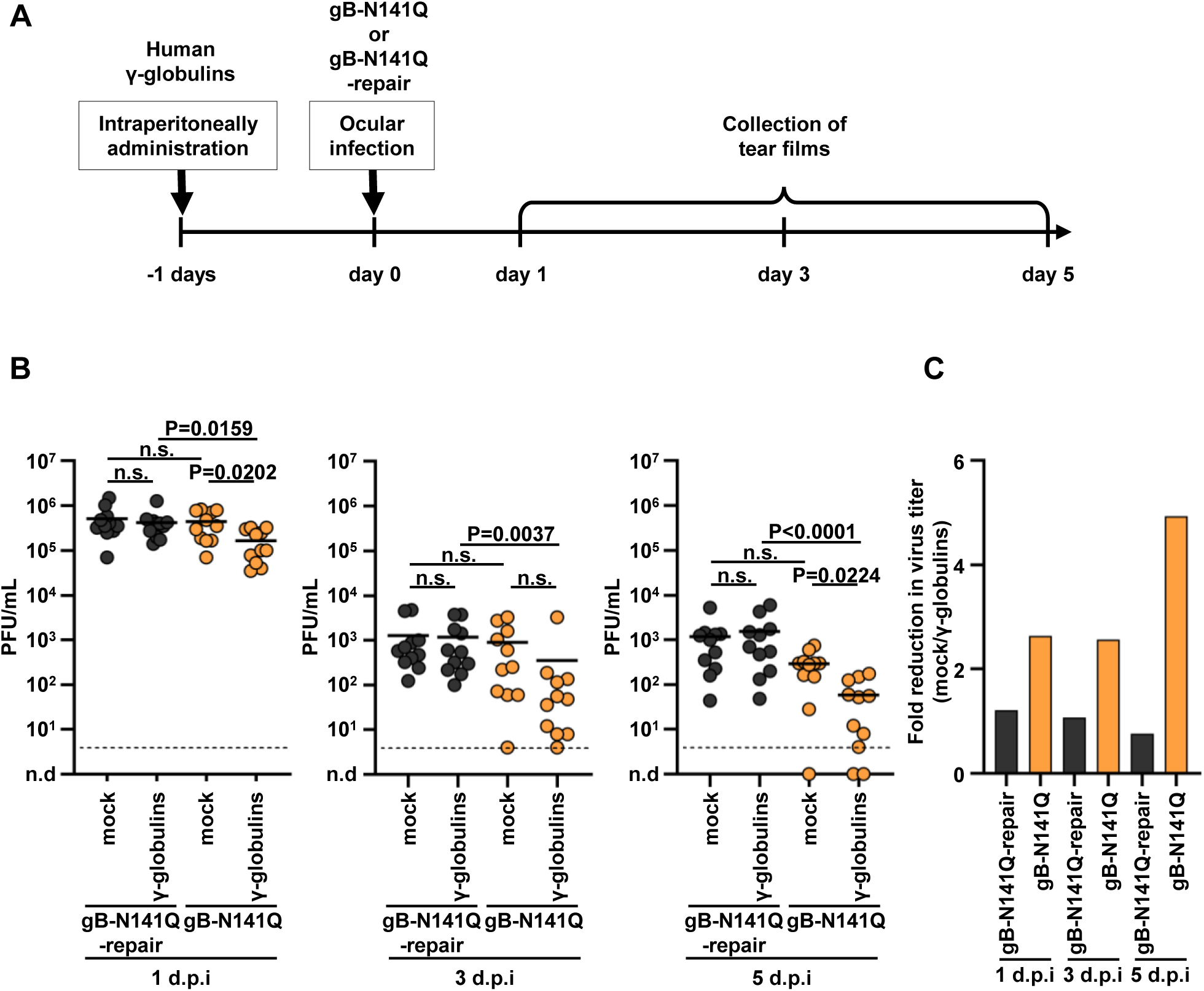
Effect of N-glycan shield at Asn-141 in gB on HSV-1 replication in the eyes of mice in the presence of human γ-globulins. (A) Schematic diagram of the experiment over time. Eleven 4-week-old female mice were mock-injected or injected intraperitoneally with human γ-globulins, and 1 day later were then ocularly infected with 3 × 10^6^ PFU/eye of gB-N141Q or gB-N141Q-repair. Samples of tear films were collected at 1, 3, and 5 days postinfection, and viral titers of the tear films were determined. (B) Viral titers of samples collected at each time point. Each data point represents the viral titer of a tear film sample from a single mouse. Horizontal bars indicate the mean for each group. Statistical analysis was performed by one-way ANOVA followed by the Tukey multiple comparisons test. n.s., not significant; d.p.i., days post infection. Dashed lines indicate limit of detection. n.d., not detected. (C) Fold reduction in mean values of viral titers due to administered human γ-globulins, which are shown in B.

### Effects of the N-glycan on gB Asn-141 on the replication of HSV-1 in the eyes of mice immunized against HSV-1

To examine the effects of the N-glycan at gB Asn-141 on HSV-1 replication in vivo in the presence of physiologically induced immunity against HSV-1, mice were subcutaneously mock-immunized or immunized with wild-type HSV-1(F). At 9 weeks after inoculation, the immunized mice were ocularly infected with gB-N141Q or gB-N141Q-repair (Fig. 7A). Samples of tear films were collected at the indicated times and viral titers of the tear films were determined (Fig. 7B). As shown in Fig. 7B, whereas the viral titers of the tear films of immunized mice infected with gB-N141Q at 1 and 2 days postinfection were comparable to those in mock-immunized mice, the viral titers of the tear films of immunized mice infected with gB-N141Q at 3 days postinfection were significantly lower than those in the mock-immunized mice. In contrast, the viral titers of the tear films of immunized mice infected with gB-N141Q-repair at 1, 2, and 3 days postinfection were comparable to those in mock-immunized mice infected with gB-141Q-repair (Fig. 7B). Thus, the ratios of gB-N141Q titers in mock-immunized mice to those in immunized mice at 3 days postinfection were higher than the ratios of gB-N141Q-repair titers in mock-immunized mice to those in immunized mice (Fig. 7C). Furthermore, the viral titers of the tear films in mock-immunized or immunized mice infected with gB-N141Q at 1 and 2 days postinfection were comparable to those in mock-immunized or immunized mice infected with gB-N141Q-repair (Fig. 7B). In contrast, the viral titers of the tear films in immunized mice infected with gB-N141Q at 3 days postinfection were significantly lower than those in immunized mice infected with gB-N141Q-repair although viral titers of the tear films in mock-immunized mice infected with gB-N141Q at 3 days postinfection were comparable to those in mock-immunized mice infected with gB-N141Q-repair (Fig. 7B). These results indicate that the presence of immunity against HSV-1 in mice inhibited replication of gB-N141Q more efficiently than the replication of gB-N141Q-repair and suggest that the N-glycan at gB Asn-141 was required for the efficient evasion of HSV-1 from immunity induced in mice previously immunized against HSV-1.

**Fig. 7.**
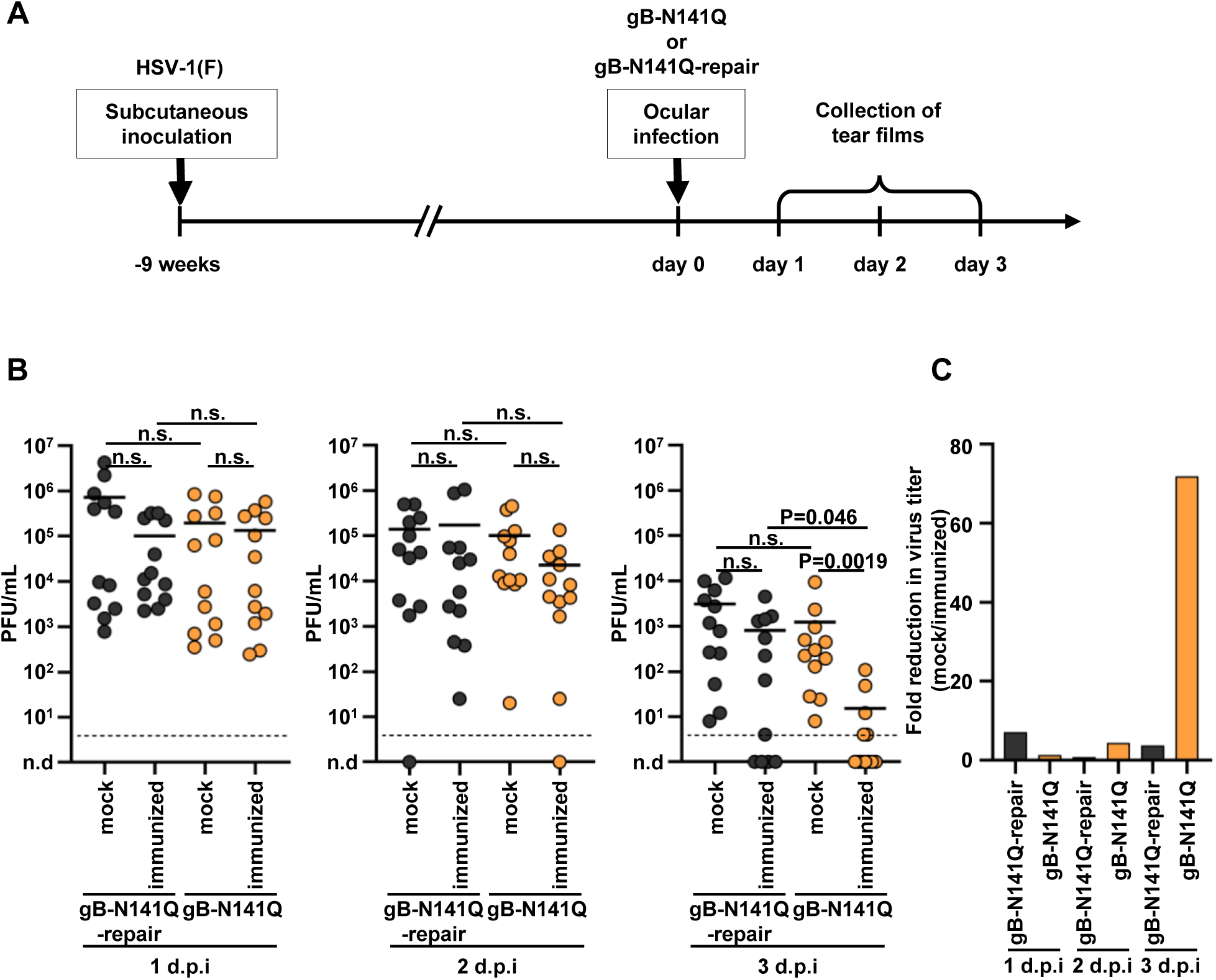
Effects of N-glycan shield at Asn-141 in gB on HSV-1 replication in the eyes of mice immunized with HSV-1. (A) Schematic diagram of the experiment over time. Twelve 3-week-old female mice were subcutaneously mock-immunized or immunized with 5 × 10^5^ PFU of wild-type HSV-1(F). At 9 weeks after inoculation, the immunized mice were ocularly infected with 3 × 10^6^ PFU/eye of gB-N141Q or gB-N141Q-repair. Samples of tear films were collected at 1, 2, and 3 days postinfection, and viral titers of the tear films were determined. (B) Viral titers of samples collected at each time point. Each data point represents the viral titer of a tear film sample from a single mouse. Horizontal bars indicate the mean for each group. Statistical analysis was performed by one-way ANOVA followed by the Tukey multiple comparisons test. n.s., not significant; d.p.i., days post infection. Dashed lines indicate limit of detection. n,d., not detected. (C) Fold reduction in mean values of viral titers due to immunization with HSV-1(F), which are shown in B.

### Effects of the N-glycan at gB Asn-141 on HSV-1 neurovirulence and replication in the CNS of naïve mice

To investigate the effects of the N-glycan at gB Asn-141 on the neurovirulence and replication of HSV-1 in the CNS of naïve mice, mice were infected intracranially with gB-N141Q or gB-N141Q-repair, and the mortality rates of these injected mice was monitored for 14 days. As shown in Fig. 8A, the mortality rate of mice infected with gB-N141Q was significantly lower than the rate of mice infected with its repaired virus (gB-N141Q-repair). We also harvested the brains of mice infected with gB-N141Q or gB-N141Q-repair at 1, 3, and 5 days postinfection and measured viral titers in their brains. As shown in Fig. 8B, the viral titers in the brains of mice infected with gB-N141Q at 1 day postinfection were comparable to those of mice infected with gB-N141Q-repair. In contrast, at later time points (3 and 5 days postinfection), viral titers in the brains of mice infected with gB-N141Q were significantly lower than the titers in the brains of mice infected with gB-N141Q-repair. These results suggest that the N-glycan at gB Asn-141 was required for efficient HSV-1 neurovirulence and replication in the CNS of naïve mice. The results also led us to investigate whether the N-glycan at gB Asn-141 acted specifically in neural cells. As shown in Fig. 8C, progeny virus yields in human neuroblastoma SK-N-SH cells infected with gB-N141Q were similar to those in the cells infected with wild-type HSV-1(F) or gB-N141Q-repair. These results further supported our observation from the results of in vitro experiments described previously that N-glycosylation on gB does not appear to play a role in the replication of HSV-1 in cell cultures.

**Fig. 8.**
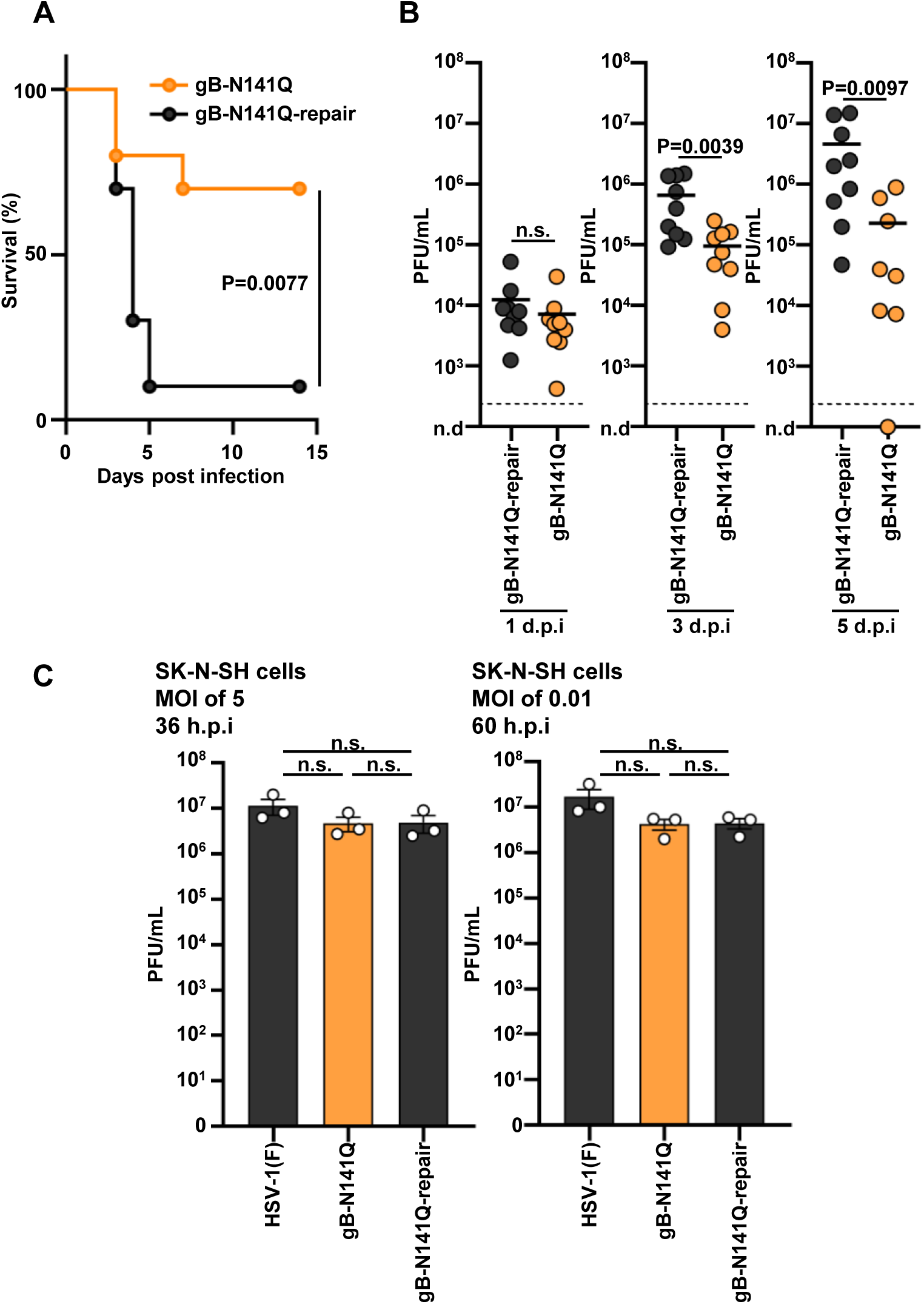
Effects of N-glycan shield at Asn-141 in gB on the neurovirulence of HSV-1 and its replication in the CNS of naïve mice. (A) Ten 3-week-old female mice were inoculated intracranially with 1 × 10^3^ PFU of gB-N141Q or gB-N141Q-repair. Infected mice were monitored for 14 days. (B) Three-week-old female mice were inoculated intracranially with 1 × 10^3^ PFU of gB-N141Q (n = 26) or gB-N141Q-repair (n = 27). At days 1 (n=9), 3 (n=9), and 5 (gB-N141Q, n=8; gB-N141Q-repair, n=9), mice from each inoculated group were sacrificed, and the viral titers in the brains were determined. Each data point is the viral titer in the brain of a single mouse. Dashed line indicates the limit of detection. n.d., not detected; d.p.i., days post infection. (C) SK-N-SH cells were infected with wild-type HSV-1(F), gB-N141Q, or gB-N141Q-repair at an MOI of 5 or 0.01. Total viruses from cell culture supernatants and infected cells were harvested at 36 h or 60 h post-infection (p.i.) and assayed on Vero cells. Each value is the mean ± standard error of the results of 3 independent experiments. Statistical analysis was performed by the log rank test (A), the two-tailed Student t test (B), or one-way ANOVA followed by the Tukey test (C). n.s., not significant.

## DISCUSSION

It is unquestionable that studies using human samples to analyze the mechanisms of infection utilized by human pathogenic viruses are crucial for understanding the mechanisms effective in humans. Considering that the gap between what can be observed from in vitro and in vivo viral infections is significant, evaluations of human samples in vivo should provide more valuable information on the mechanisms of infection than evaluations of the samples in vitro. However, there has been a lack of in vivo information because human samples that could be used for in vivo analyses as well as in vivo models that could represent the pathogenesis of viral infections in humans are limited.

In this study, we clarified a novel immune evasion mechanism used by HSV-1, namely, that a glycan shield at Asn 141 of the HSV-1 gB mediated evasion from the deleterious effects of human antibodies such as in vitro neutralization and ADCC. Our observations are supported by the molecular model of N-glycosylation at gB Asn-141, which is based on the prefusion structure of HSV-1 gB (18). The model predicts that N-glycosylation at gB Asn-141 masked 27 amino acids in the gB molecule. Of these amino acids, 70% have been mapped to the functional regions of gB, FR2, and FR3, which were previously identified according to the known epitopes of various neutralizing monoclonal antibodies (20, 21). Notably, the N-glycosylation at gB Asn-141 was predicted to mask Asp-419, a residue critical for the binding of gB to the C226 antibody, which shows high neutralizing activity in preventing the association of gB with a complex of gH and gL and fusion (21). Furthermore, the amino acids predicted to be masked by glycosylation at Asn-141 included or were positioned near those (Pro-361, Asp-408, Asp-419, Asn-430, Asn-458, Arg-470, Pro-481, Ile-495 and Thr-497) previously shown to be critical for gB receptor- or gD receptor-mediated fusion (22–24), introducing the possibility that antibodies target these amino acid residues.

Our observations that the glycan shield on HSV-1 gB seemed to significantly increase viral replication in the eyes of mice not only in the presence of HSV-1 immunity but also in the presence of human antibodies, supports our prediction from the clarified in-vitro effects of the glycan shield on HSV-1 that the glycan shield would be effective in the presence of human antibodies in vivo and probably in human beings. Notably, we also presented evidence that the glycan on HSV-1 gB was required for efficient viral neurovirulence and replication in the CNS of naïve mice. Thus, we have identified an important glycan shield on the HSV-1 gB that appears to have 2 affects: evasion from human antibodies in vivo and neurovirulence in naïve hosts. Considering the clinical features of HSV-1 infection, these results suggest that the glycan shield not only facilitates recurrent HSV-1 infections in latently infected humans by evading antibodies, but also is important for HSV-1 pathogenesis during the initial infection.

In this study, we used pooled human γ-globulins from human blood as human antibodies. HSV-1 is a ubiquitous human pathogen; approximately 70% of the global human population is infected with HSV-1, and most HSV-1-infected humans have been reported to be latently infected with the virus (1–3, 15). Therefore, it is conceivable that the effects of pooled human γ-globulins represent the effects of antibodies in humans latently infected with HSV-1. Thus, the mouse model with passive transfer of pooled human γ-globulins used in this study potentially mimicked the in vivo effects of human antibodies in humans latently infected with HSV-1. HSV-1 frequently reactivates from latent infections and is transmitted to new human hosts. Therefore, the host’s immune responses to HSV-1 persist in latently infected humans because of the repeated stimulation of the immune system, resulting in the progressive enhancement of long-term immunity (1–3). The viral strategy of using the glycan shield on HSV-1 gB to evade antibodies, which was clarified in this study, may protect reactivated viruses from existing antibodies to HSV-1 in latently infected humans and thereby facilitate their transmission to new human hosts. Notably, the N-glycosylation site on HSV-1 gB is widely conserved in viruses subclassified in the alphaherpesvirus subfamily of herpesviruses (25), suggesting that this is a general viral mechanism of evasion from the immune system. Moreover, clarification of the HSV-1 mechanism of evasion from antibodies supports earlier conclusions (5–9) that were based on previous clinical trials of HSV vaccines; namely, that antibodies are essential to the control of HSV-1 infections in humans. Additional studies to reveal other glycan shields against human antibodies on HSV envelope glycoproteins are important and should be of interest. Those studies and this present study may provide insights into the design of effective therapeutic HSV vaccines against frequent recurrences of herpes virus infections such as genital herpes.

Previous studies have characterized N-glycosylation at Asn-133 of the HSV-2 gB and at Asn-154 of the pseudorabies virus (PRV) gB, which correspond to the N-glycosylation at Asn-141 of HSV-1 gB (26, 27). None of those studies addressed the effects of the N-glycosylation of those viruses’ gB on evasion from antibodies and pathogenesis in vivo. In agreement with our observation in this study that the N141Q mutation in HSV-1 gB did not affect viral replication in Vero and SK-N-SH cells, the ectopic expression of the PRV gB-N154Q mutant rescued the entry deficiency of a gB-deficient PRV so that its entry would be at a level similar to that of PRVs with wild-type gB (27). In contrast, the ectopic expression of the HSV-2 gB-N133Q mutant barely rescued the entry deficiency of a gB-deficient HSV-2 (26). As observed with the N141Q mutation in gB of HSV-1 in the context of viral infection, the ectopic expression of both the HSV-2 gB-N133Q and PRV gB-N154Q mutant showed impaired cell surface expression of the mutants. These observations point out both the similarities in and differences between the roles of the glycosylation of gB in viruses.

## ACKNOWLEDGEMENTS

We thank Risa Abe, Keiko Sato, Tohru Ikegami, and Yui Muto for their excellent technical assistance. We are grateful to Seiya Yamayoshi for helpful discussions with the ADCC assays. This study was supported by Grants for Scientific Research and Grant-in-Aid for Scientific Research (S) (20H05692) from the Japan Society for the Promotion of Science (JSPS), grants for Scientific Research on Innovative Areas (21H00338, 21H00417, 22H04803) and a grant for Transformative Research Areas (22H05584) from the Ministry of Education, Culture, Science, Sports and Technology of Japan, a PRESTO grant (JPMJPR22R5) from Japan Science and Technology Agency (JST), grants (JP20wm0125002, JP20wm0225009, JP20wm0225017, JP22fk0108640, JP22gm1610008, JP223fa627001) from the Japan Agency for Medical Research and Development (AMED), grants from the International Joint Research Project of the Institute of Medical Science, the University of Tokyo, and grants from the Takeda Science Foundation, the Uehara Memorial Foundation and the Mitsubishi Foundation.

## MATERIALS AND METHODS

### Cells and Viruses

Vero, HEK293FT, Plat-GP, and SK-N-SH cells were described previously (28, 29). Wild-type HSV-1(F) was described previously (30).

### Plasmids

The construction of pFLAG-CMV2-EGFP and pcDNA-MEF-gB was described previously (28, 31). First, to construct pBS-KanR-ePheS*, oligonucleotides making up a kanamycin resistant (KanR) cassette with the I-SceI recognition site and an ePheS* cassette, which has T251A/A294G mutations in ePheS, were amplified from the DNA templates of pEPkan-S (13) and pUC18K ePAG2 (14), respectively. The primers that were used are listed in S-Table 1B. Then, the two linear DNA fragments were fused by PCR and cloned into the pBluescript KS(+) (Stratagene), as described previously (32). Second, pBS-TEV-2xStrep-KanS was constructed by cloning the kanamycin resistance (KanR) cassette with the I-SceI recognition site, which was amplified by PCR from the pEPkan-S template with the use of primers that additionally encoded the Tobacco Etch Virus (TEV) protease cleavage-site and tandem-strep epitopes as listed in S-Table 1B, into the pBluescript KS(+). Third, pcDNA3.1-tagRFP-P2A and pcDNA3.1-P2A-tagRFP were constructed by cloning the tagRFP open reading frame (ORF), which was amplified by PCR from ptagRFP-N1 (33) with the use of primers that additionally encoded the 2A self-cleaving peptide fused to its carboxyl terminus (tagRFP-P2A) or amino-terminus (P2A-tagRFP) as listed in S-Table 1B, respectively, into pcDNA3.1 (Invitrogen). Fourth, pcDNA3.1-tagRFP-P2A-stop was constructed by cloning annealed DNA oligonucleotides listed in S-Table 1B into pcDNA3.1-tagRFP-P2A. Fifth, pcDNA3.1-gB-P2A-tagRFP was constructed by cloning gB ORF, which was amplified by PCR from the HSV-1(F) genome isolated as described previously (34), using the primers listed in S-Table 1B into pcDNA3.1-P2A-tagRFP by the In-Fusion HD Cloning Kit (Takara), according to the manufacturer’s instructions. Sixth, pcDNA3.1-gD-P2A-tagRFP was constructed by cloning gD ORF, which was amplified by PCR from the HSV-1(F) genome using primers listed in S-Table 1B into pcDNA3.1-P2A-tagRFP by the In-Fusion HD Cloning Kit.

To construct pRetroX-TRE3G-gBo and pRetroX-TRE3G-ICP4o, the sequences of codon-optimized UL27(gBo) and α4 (ICP4o), which are shown in S-Table 1C, were engineered according to the GenScript’s OptimumGene algorithm, and then synthesized and cloned into pRetroX-TRE3G (Takara) by GenScript.

### Establishment of stable Vero cells with tetracycline-inducible codon-optimized gB and ICP4 (gBo and ICP4o) expression

Vero cells were transduced with supernatants of Plat-GP cells cotransfected with pMDG (35) and pRetroX-Tet3G (TaKaRa), selected with 1 mg/mL G418 solution (Wako) to generate Tet3G-Vero cells. The cells were further transduced with a mixture of supernatants of Plat-GP cells cotransfected with pMDG and pRetroX-TRE3G-gBo, and supernatants of Plat-GP cells co-transfected with pMDG and pRetroX-TRE3G-ICP4o to establish gBo/ICP4o-TetON-Vero cells. After double selection with 1 mg/mL of G418 solution and 5μg/mL of puromycin, a single clone in which expression of gBo and ICP4o was induced by doxycycline was selected.

### Two-step Red-mediated recombination using the KanR/ePheS* cassette

The two-step Red-mediated mutagenesis procedure used in this study was performed as described previously (13, 36). Briefly, linear DNA fragments containing an I-SceI recognition sequence, KanR and ePheS*cassettes, and target homologous sequences were amplified by PCR from pBS-KanR-ePheS* using the primers listed in S-Table 1D. The linear fragments were electroporated into the electrocompetent *Escherichia coli* strain GS1783 containing the pYEbac102Cre (30, 37). The transformed bacteria were then incubated at 32℃ for 40 to 60 min and plated on LB agar plates containing 20 μg/mL of chloramphenicol and 40 μg/mL of kanamycin to select *E. coli* clones harboring pYEbac102Cre containing the KanR and ePheS* cassettes (KanR/ePheS* cassettes). Kanamycin-resistant colonies were screened by PCR with the appropriate primers. Next, the KanR/ePheS* cassettes were excised by expressing the I-SceI homing enzyme in GS1783 through induction with arabinose, followed by induction of the Red recombination machinery by raising the temperature. Briefly, 100 μL of an overnight culture of kanamycin-resistant *E. coli* clones grown in LB medium containing chloramphenicol and kanamycin was inoculated into 2 mL of LB medium containing chloramphenicol only. Bacteria were incubated at 32°C for 2 to 4 h with shaking, followed by addition of 10% (wt/vol) L-arabinose (Wako) to the culture at a 1:5 ratio, and incubated for another 1 h at 32°C. Finally, the *E. coli* culture was incubated at 42°C for 30 min. It was then shaken at 32°C for another 1 to 2 h, and 50 μL of 10^-3^ to 10^-4^ dilutions of the culture were plated onto LB agar plates containing 20 μg/mL of chloramphenicol and 1 mM of 4-chloro-phenylalanine (4CP) to select *E. coli* clones harboring the pYEbac102Cre, from which the KanR/ePheS* cassette was excised. Chloramphenicol- and 4CP-resistant colonies were screened by PCR with appropriate primers, which was followed by nucleotide sequencing for confirmation of the desired mutation.

### Generation of recombinant HSV-1

Recombinant viruses YK650 (UL51-T190A_KanR/ePheS*), YK681 (gB-N87Q), YK683 (gB-N141Q), YK685 (gB-N398Q), YK687 (gB-N430Q), YK689 (gB-N489Q), YK691 (gB-N674Q), YK693 (gB-N888Q), YK682 (gB-N87Q-repair), YK684 (gB-N141Q-repair), YK686 (gB-N398Q-repair), YK688 (gB-N430Q-repair), YK690 (gB-N489Q-repair), YK692 (gB-N674Q-repair), and YK694 (gB-N888Q-repair) (Fig. 1) were generated by the two-step Red-mediated mutagenesis procedure using the KanR/ePheS* cassette as described in the previous section with the primers listed in S-Table 1D. The recombinant virus YK649 (UL51-T190A_KanR) was generated by the two-step Red-mediated mutagenesis procedure using *E. coli* GS1783 containing pYEbac102Cre, as described previously (13, 36), with the exception that the primers used instead of those described previously are listed in S-Table 1D. The recombinant virus YK695 (ΔgB), in which the UL27 gene encoding gB was disrupted by deleting gB codons 1-727 with a kanamycin resistance gene, was generated by the two-step Red-mediated mutagenesis procedure using *E. coli* GS1783 containing pYEbac102Cre, as described previously (13, 36), with the exception that the primers used instead of those described previously are listed in S-Table 1D.

The recombinant virus YK696 (ΔgB-repair), in which the deletion mutation in gB was repaired, was generated by cotransfection with pYEbac102Cre carrying the gB-deletion mutation and pCRxgB (38) into Vero cells. Plaques were isolated and purified on Vero cells. Restoration was confirmed by nucleotide sequencing.

The recombinant virus YK717 (gB-SE), which expresses gB fused to a TEV protease cleavage site and a Strep-tag; and recombinant virus YK718 (gD-SE), which expresses gD fused to a TEV protease cleavage site and a Strep-tag, were generated by the two-step Red-mediated mutagenesis procedure using *E. coli* GS1783 containing pYEbac102Cre, as described previously (13, 36), with the exception that the primers used instead of those described previously are listed in S-Table 1D.

In experiments in which YK695 (HSV-1 ΔgB) was used, viruses were propagated and assayed in HSV-1 gBo/ICP4o-TetON-Vero cells in the presence of doxycycline (DOX) (1 mg/mL). Other viruses used in this study were propagated and titrated in Vero cells.

### Antibodies

Commercial antibodies used in this study were mouse monoclonal antibodies to gB (H1817; Virusys) and α-tubulin (DM1A; Sigma), and rabbit polyclonal antibodies to VP23 (CAC-CT-HSV-UL18; Cosmo Bio).

### PNGase F Digestion and immunoblotting

Vero cells were infected with each of the indicated viruses at an MOI of 5 for 24 h and lysed with T-PER Tissue Protein Extraction Reagent (Thermo Scientific). The lysates were sonicated and denatured with Glycoprotein Denaturing Buffer (NEB) by heating them at 100°C for 10 minutes. Aliquots of the lysates were incubated with 2500 units of PNGase F (NEB) at 37°C for 1 h. Aliquots of the lysates incubated under the same conditions without PNGase F were used as controls. The incubated mixtures were subjected to immunoblotting as described previously (39).

### Detection of gB- or gD-specific antibodies in pooled human γ-globulins by flow cytometry

PEI MAX (Polyscience, Inc.) was used to transfect HEK293FT cells with selected plasmids. At 48 h post-transfection, the transfected cells were detached from their culture plates and washed once with PBS supplemented with 2% FCS (washing buffer). Cells were fixed and permeabilized with Cytofix/Cytoperm (Beckton Dickinson) and incubated with diluted human γ-globulins (G4386; Sigma) on ice for 30 min. After the cells were washed with washing buffer, they were further incubated with anti-human IgG conjugated to Alexa Flour 647 (Invitrogen) on ice for 30 min. After the cells were washed again, they were analyzed with a CytoFLEX S flow cytometer (Beckman Coulter). The data were analyzed with FlowJo 10.8.1 software (Becton Dickinson).

### Depletion of gB- or gD-specific antibodies from pooled human γ-globulins

Vero cells (5 × 10^7^) were infected with HSV-1(F), gB-SE, or gD-SE at an MOI of 0.5 for 24 h and lysed in 5 mL of radioimmunoprecipitation assay (RIPA) buffer (10 mM Tris-HCl [pH 7.4], 150 mM NaCl, 1% Nonidet P-40 [NP40], 0.1% deoxycholate, 0.1% sodium deodecyl sulfate, 1 mM EDTA) containing a protease inhibitor cocktail (Nacalai Tesque). After centrifugation, the supernatants were precleared by incubating with protein G-Sepharose beads (GE Healthcare), and reacted with 100 μL of Strep-Tactin Sepharose beads (IBA Lifesciences) for 4 h at 4°C. The beads were collected by brief centrifugation and washed 4 times with RIPA buffer and 2 times with PBS. Samples of gB-SE or gD-SE immobilized on Strep-Tactin Sepharose beads were incubated with 1 mL of diluted human γ-globulins (G4386; Sigma) (0.082 and 3.04 mg/mL in medium 199 containing 1% FCS and ADCC assay buffer for neutralization assays and ADCC assays, respectively) at 4°C overnight; and after centrifugation, the supernatants containing gB- or gD-antibody depleted human γ-globulins were filtered.

Similarly, HSV-1(F)-infected Vero cell lysates prepared as described for the depleted human γ-globulins was incubated with Strep-Tactin Sepharose beads, and after centrifugation and washing, the beads were incubated with human γ-globulins diluted as described, and then centrifuged, followed by filtration of the supernatants to produce samples of mock-depleted human γ-globulins.

### Neutralization assay

Pooled human γ-globulins (G4386; Sigma) serially diluted in medium 199 containing 1% FCS were mixed 1:1 with 100 PFU of each selected virus in medium 199 containing 1% FCS, incubated at 37°C for 1 h, and then inoculated onto Vero cell monolayers to perform plaque assays. At 2 days postinfection, the plaques were counted. The percentage of neutralization was determined as follows: the numbers of plaques formed by the virus samples that had been incubated with or without human γ-globulins as a value with the following formula: 100 × [1-(numbers of plaques produced after incubation of viral samples with human γ-globulins)/(numbers of plaques produced after incubation of viral samples without human γ-globulins)].

### ADCC reporter assay

The extent of ADCC activation induced by human γ-globulins was evaluated with the use of an ADCC Reporter Bioassay (Core Kit; Promega, G7010) and the EnSpireMultimode Plate Reader (PerkinElmer). The assay was used according to the manufacturer’s instructions. Briefly, Vero cells were infected at an MOI of 1 with each selected virus in medium 199 containing 1% FCS. After adsorption for 1 h, the inoculum was removed, and the cell monolayers were overlaid with medium 199 containing 10% FCS. At 24 h post-infection, the culture medium was replaced with ADCC assay buffer containing effector cells and diluted human γ-globulins at a 2:1 ratio and incubated at 37°C for 6 h. Bio-Glo luciferase reagent was then added, and the luciferase signals were quantitated as relative light units (RLUs) on an EnSpire reader. Extent of induction was calculated as follows: Fold induction = (RLUs_with antibody_ - RLUs_background_)/(RLUs_no antibody_ - RLUs_background_).

### Determination of gB expression on the surfaces of HSV-1-infected cells

The expression of HSV-1 glycoproteins on the surfaces of infected cells was analyzed as described previously (40). Briefly, Vero cells were infected at an MOI of 1 with each selected virus in medium 199 containing 1% FCS. After adsorption for 1 h, the inoculum was removed, and the cell monolayers were overlaid with medium 199 containing 10% FCS. At 24 h post-infection, cell monolayers were detached with PBS containing 0.02% EDTA and were washed 1 time with PBS supplemented with 2% FCS (washing buffer). To analyze the total expression of gB, infected Vero cells were detached as described, fixed, and permeabilized with Cytofix/Cytoperm Fixation/Permeabilization Solution (Becton Dickinson). Treated and untreated cells were then incubated with mouse anti-gB monoclonal antibody in washing buffer on ice for 30 min. After the cells were washed with washing buffer, they were further incubated with anti-mouse IgG conjugated to Alexa Flour 647 dye (Invitrogen) on ice for 30 min. After the cells were washed again, they were analyzed with a CytoFLEX S flow cytometer (Beckman Coulter). The data were analyzed by FlowJo 10.8.1 software (Becton Dickinson).

### Immunofluorescence assays

Immunofluorescence assays were performed as described previously (41).

### Animal studies

Female ICR mice were purchased from Charles River Laboratories. For ocular infections by each selected virus in mice in the presence of pooled human γ-globulins, four-week-old mice were injected intraperitoneally with 1250 mg/kg of human γ-globulins or PBS. One day after administration, the mice were ocularly infected with 3 × 10^6^ PFU/eye of each selected virus, as described previously (29). For mice immunized with HSV-1 before ocular infection, three-week-old mice were injected subcutaneously in the neck with 5 × 10^5^ PFU of HSV-1(F). The immunized mice were then infected ocularly 9 weeks after immunization with 3 × 10^6^ PFU/eye of each selected virus as described previously (29). Virus titers in the tear films of mice were determined as described previously (42).

For intracranial infections, three-week-old mice were inoculated intracranially with each selected virus as described previously (29). Mice were monitored daily, and mortality occurring from 1 to 14 days postinfection was attributed to the infecting virus. To measure viral titers in the brains of infected mice, three-week-old female ICR mice were each inoculated intracranially with 1 × 10^3^ PFU of each selected virus. At 1, 3, and 5 days postinfection, the brains of the mice were harvested, and virus titers were determined on Vero cells. All animal experiments were carried out in accordance with the Guidelines for Proper Conduct of Animal Experiments, Science Council of Japan. The protocol was approved by the Institutional Animal Care and Use Committee (IACUC) of the Institute of Medical Science, The University of Tokyo (IACUC protocol approval number: A21-55).

### Modeling of the N-glycosylated gB protein

The N-glycan core (Man_3_GlcNAc_2_) was modeled according to the prefusion structure of HSV-1 (18) gB (PDB ID: 6Z9M) using the Glycan Reader and Modeler (43) and the CHARMM-GUI program (44, 45). Discovery Studio 2021 software (Dassault Systèmes) was used to change the χ1 angle (N-Cα-Cβ-Cγ) of the Asn141 B chain (PDB ID: 6Z9M) from -178° to -66° to avoid steric clash of the N-glycan with the neighboring polypeptide. Visualization of the protein 3D structure was performed in the PyMol Molecular Graphics System, version 2.5 (Schrödinger, LLC).

### Analysis of protein surface

The AREAIMOL program (CCP4 package, version 6.2) was used to determine the accessible surface area (ASA) (46). A spherical probe with a radius of 10 Å, which is similar to the dimension of the antigen-binding fragments (single-chain variable fragment [scFv]) of the antibodies, was used in the estimation of the ASA (19, 47). The extent of glycan shielding (ΔASA) was estimated for each amino acid residue by calculating the difference between the ASAs of N-glycosylated and nonglycosylated gB structures (ΔASA = ASA [nonglycosylated gB] – ASA [N-glycosylated gB]).

### Statistical analysis

The unpaired t test was used to compare 2 groups. One-way or two-way ANOVA followed by the Tukey or Dunnett multiple comparisons tests were used for multiple comparisons. A *P* value < 0.05 was considered significant. For the statistical analysis of viral titers, data were converted to common logarithms (log_10_). For values below the detection limit, statistical processing was performed assuming that the values are those of the detection limit. GraphPad Prism 8 (GraphPad Software) was used to perform statistical analysis.

---

**Supplementary Fig. 1.**
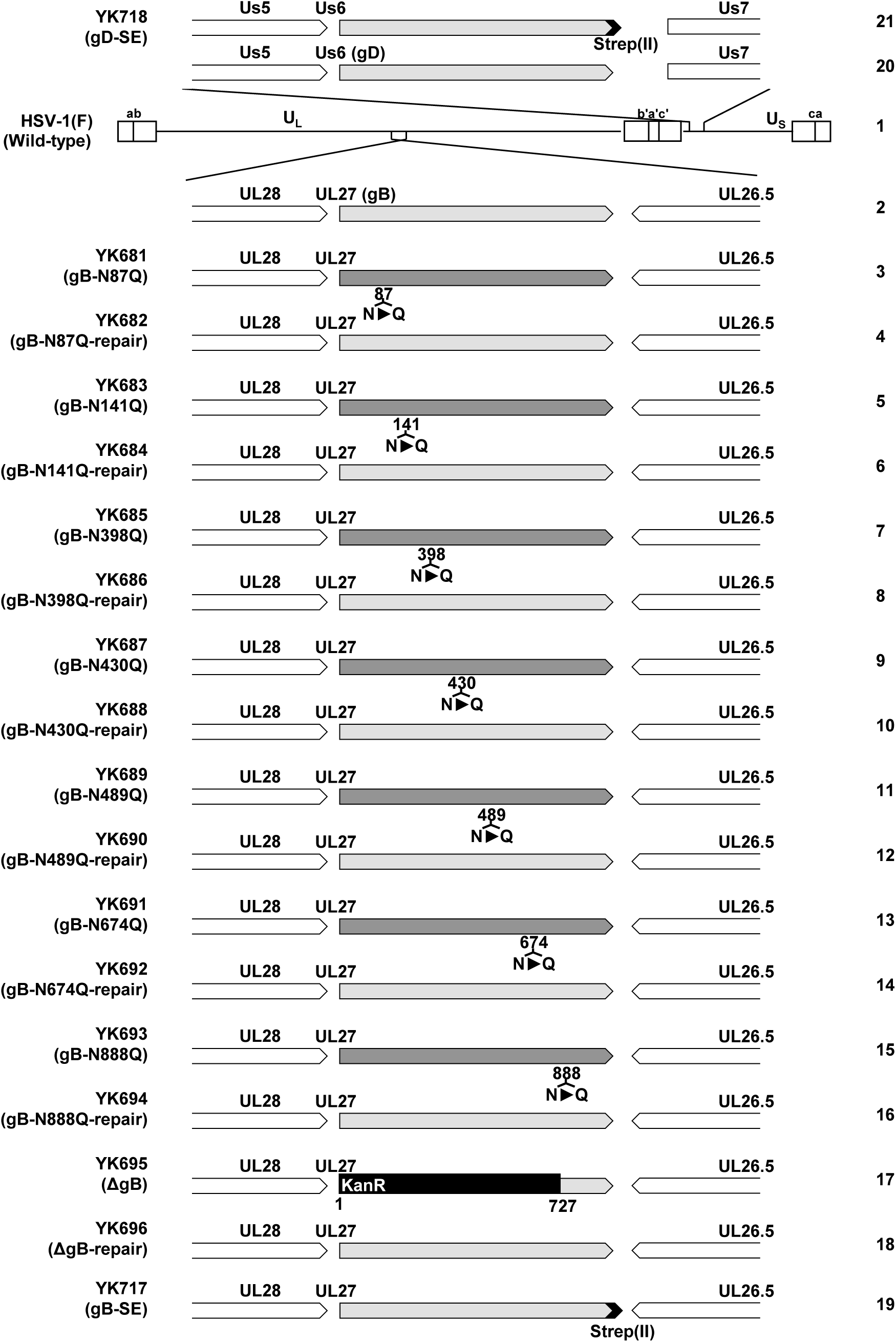
Schematic diagrams of wild-type HSV-1(F) and the creation of its recombinants that were used in this study. Schematic diagrams of the genomic structures of wild-type HSV-1(F) and the recombinant viruses used in this study. Line 1, wild-type HSV-1(F) genome; line 2, domains of the UL26.5 to UL28 genes; lines 3 to 19, recombinant HSV-1 with mutations in the UL27 gene encoding gB; line 20, domains of the Us5 to Us7 genes; line 21, recombinant HSV-1 with mutation in the Us6 gene encoding gD.

**Supplementary Fig. 2.**
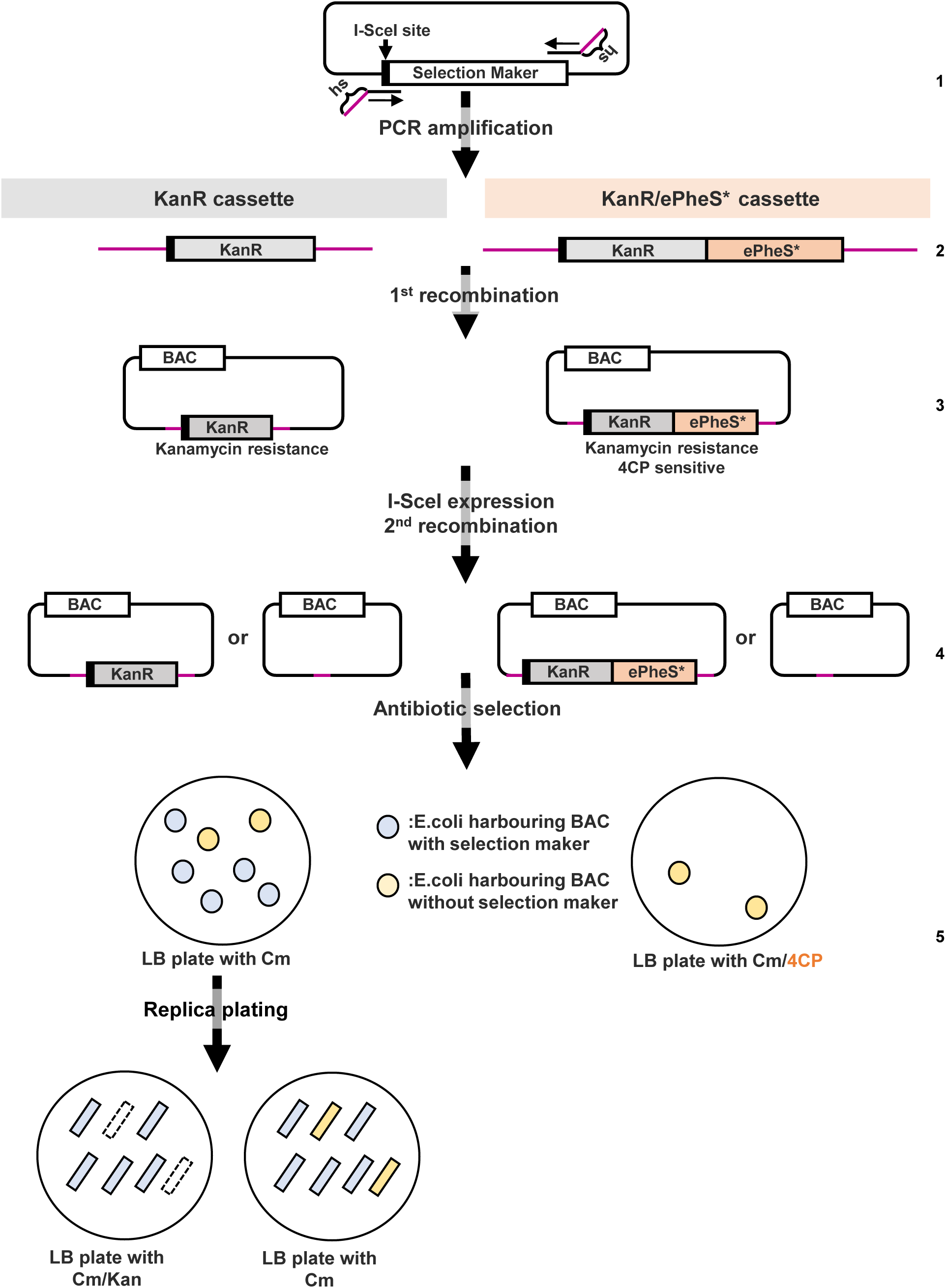
Flow chart of two-step Red-mediated recombination using the KanR or KanR/ePheS* cassette. Line 1, plasmid with a selection marker and an I-SceI site used as a template for PCR amplification; Line 2, PCR-amplified linear DNA fragments containing target homologous sequences (hs) and KanR or KanR/ePheS* cassette; Line 3, target BAC clones in which KanR or KanR/ePheS* cassette was inserted by Red recombination (1^st^ recombination); Line 4, KanR or KanR/ePheS* cassette was excised by expression of the I-SceI restriction enzyme, followed by the second Red recombination (2^nd^ recombination). In this step, the *E. coli* clones harboring BAC in which the KanR or KanR/ePheS* cassette were not removed remained; Line 5, selection of *E. coli* clones harboring the BAC in which the KanR or KanR/ePheS* cassette was removed. When the KanR cassette was used, the *E. coli* clones were selected by replica plating using agar plates containing chloramphenicol (Cm) or agar plates containing Cm and kanamycin (Kan). When the KanR/ePheS* cassette was used, the *E. coli* clones were selected without replica plating and with agar plates containing Cm and 4-chloro-phenylalanine (4CP).

**Supplementary Fig. 3.**
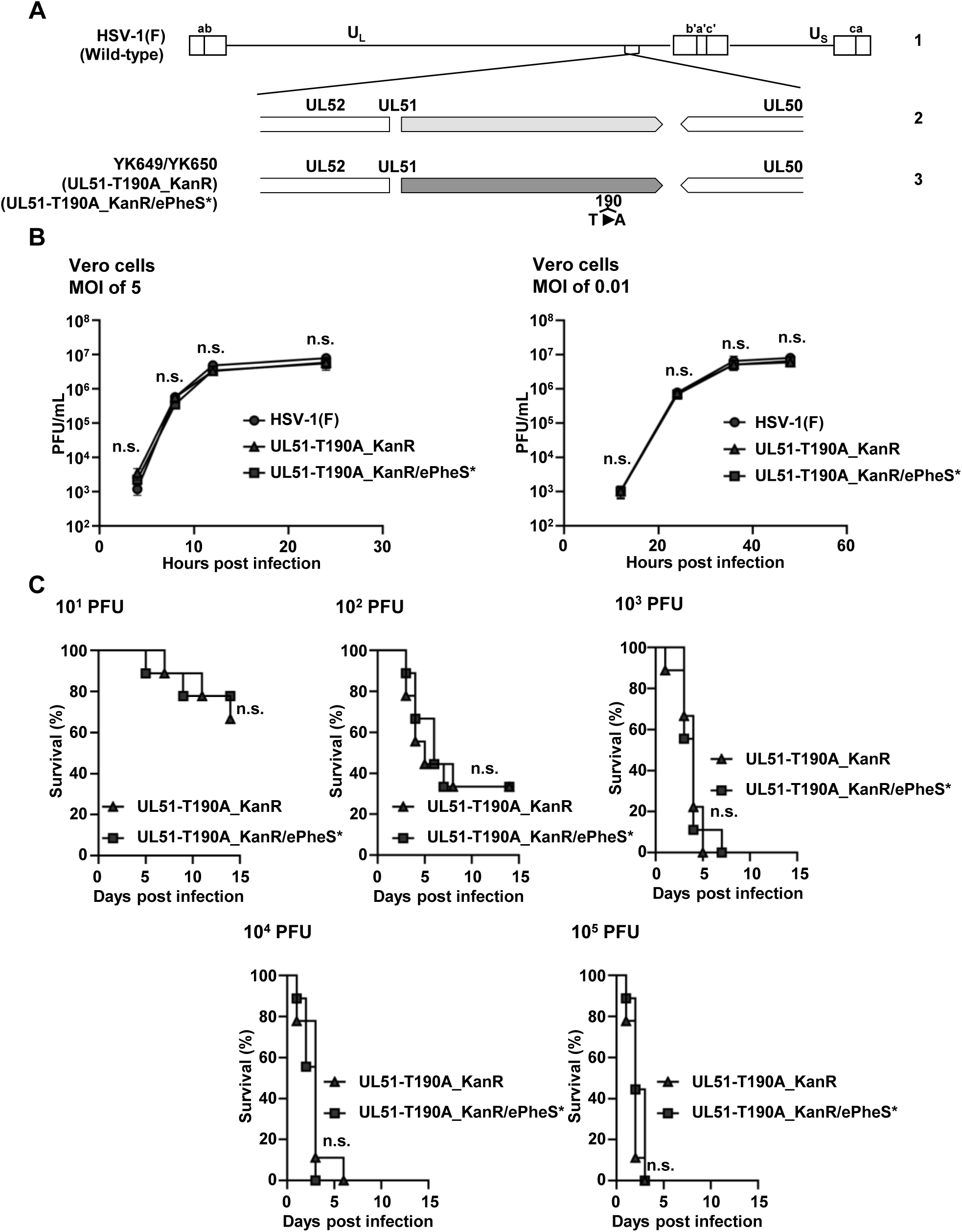
Characterization of recombinant HSV-1 generated with the use of the improved HSV-1 genetic manipulation system. (A) Schematic diagrams of recombinants of HSV-1 used in these experiments. Recombinant viruses YK649 (UL51-T190A_KanR) and YK650 (UL51-T190A_KanR/ePheS*) were generated by the original and improved HSV-1 genetic manipulation systems, respectively. (B) Vero cells were infected at an MOI of 5 or 0.01 with wild-type HSV-1(F), YK649 (HSV-1 UL51-T190A_KanR), or YK650 (HSV-1 UL51-T190A_KanR/ePheS*). All the recombinant viruses from each cell culture supernatant plus the infected cells were harvested at the indicated times, and a sample of each harvested recombinant was assayed on Vero cells. Each value is the mean ± standard error of the results of 3 independent experiments. Statistical analysis was performed by one-way ANOVA followed by the Tukey test. n.s., not statistically significant between HSV-1(F) and YK649 (UL51-T190A_KanR), HSV-1(F) and YK650 (UL51-T190A_KanR/ePheS*), and YK649 (UL51-T190A_KanR) and YK650 (UL51-T190A_KanR/ePheS*). (C) Nine 3-week-old female ICR mice were infected intracranially with the indicated amounts of YK650 (UL51-T190A_KanR/ePheS*) or YK649 (UL51-T190A_KanR), and monitored for 14 days. Statistical analysis was performed by the log rank test. n.s., not significant.

**Supplementary Fig. 4.**
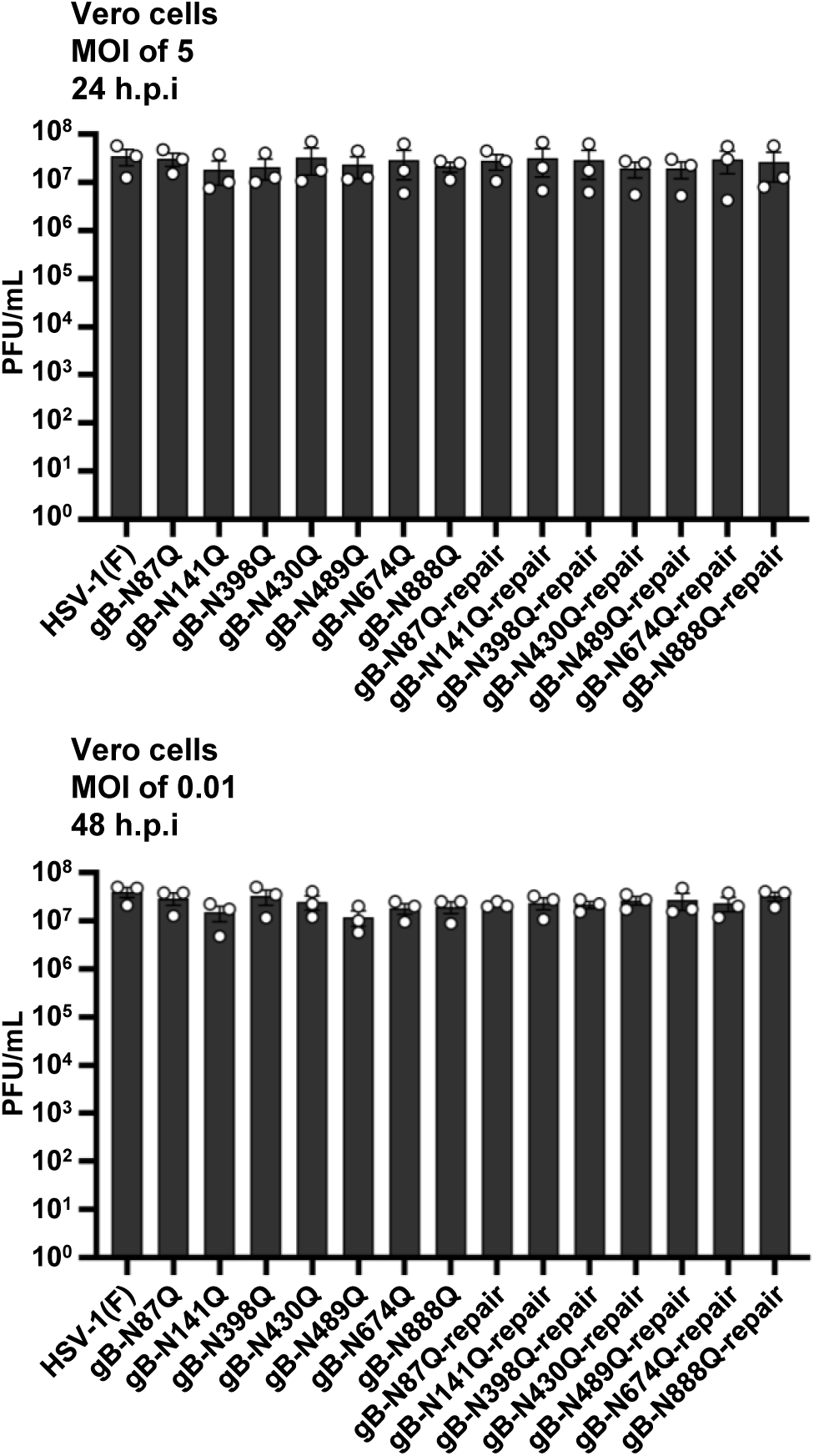
Effect of mutation in each of the gB N-glycosylation sites on HSV-1 replication in cell cultures. Vero cells were infected with wild-type HSV-1(F), each of the gB mutant viruses, or each of their repaired viruses at an MOI of 5 or 0.01. Total virus from cell culture supernatants and infected cells was harvested at 24 or 48 h postinfection and assayed on Vero cells. Each value is the mean ± standard error of the results from 3 independent experiments. There were no statistically significant differences between the amounts of any mutant virus compared to the amount of HSV-1(F) by one-way ANOVA followed by the Dunnett test.

**Supplementary Fig. 5.**
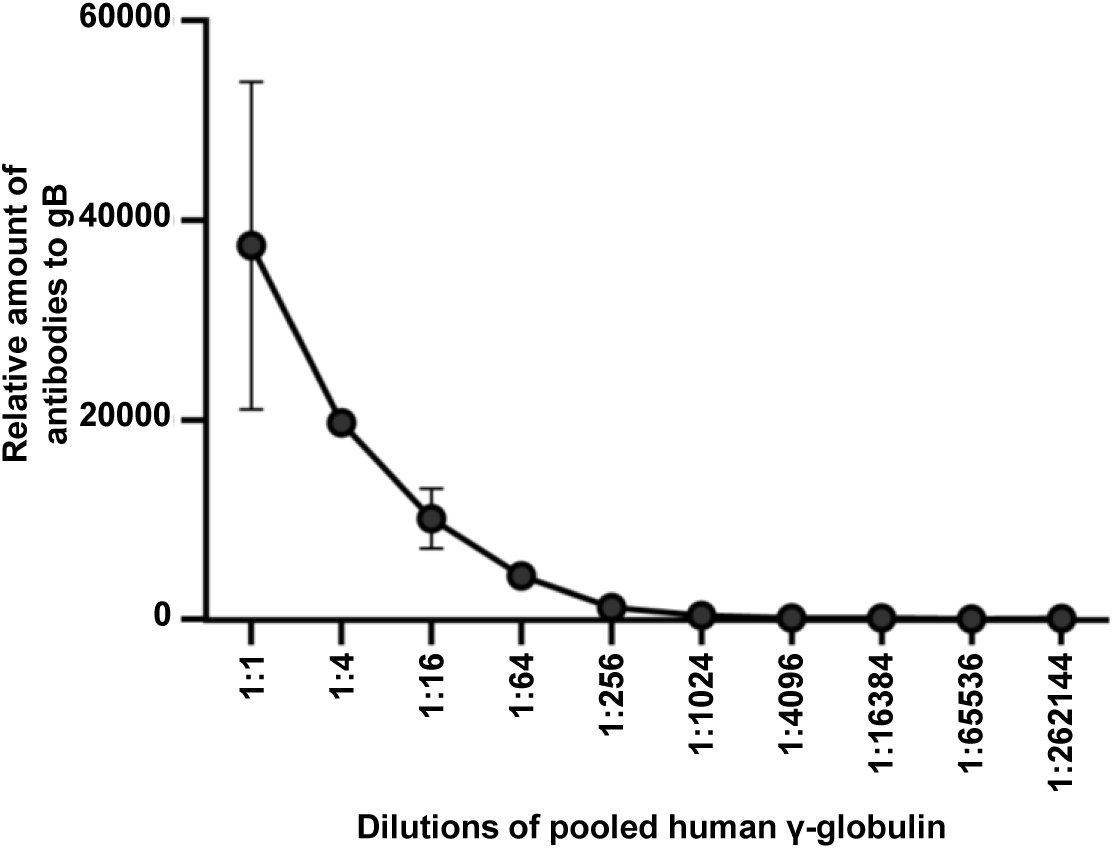
Detection of gB antibodies in human γ-globulins. HEK293FT cells were transfected with pFLAG-CMV2-EGFP or pcDNA-MEF-gB. Transfected cells were fixed, permeabilized, and incubated with serially diluted human γ-globulins (1.31 mg/mL) and analyzed by flow cytometry. The relative amounts of gB antibodies were calculated as follows: (mean fluorescent intensity of cells transfected with pcDNA-MEF-gB) − (mean fluorescent intensity of cells transfected with pFLAG-CMV2-EGFP). Data are means ± standard error of the results of 2 independent experiments.

**Supplementary Fig. 6.**
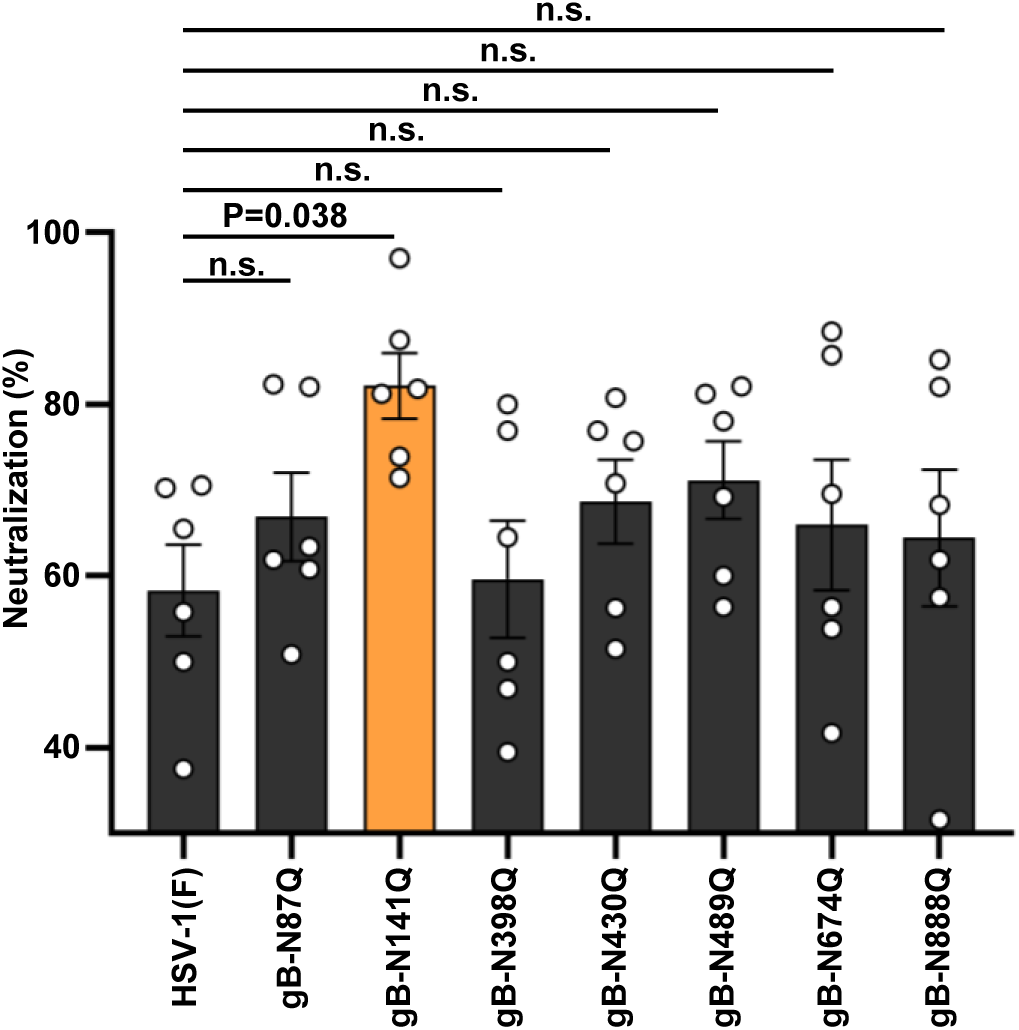
Effect of mutation in each of the gB N-glycosylation sites on viral susceptibility to neutralization by human γ-globulins. 100 PFU of wild-type HSV-1(F) or each of the gB mutant viruses were incubated with 0.041 mg/mL human γ-globulins at 37°C for 1 h, and then inoculated onto Vero cell monolayers for plaque assays. The percentage of neutralization was calculated from the number of plaques formed by each of the viruses that were incubated with or without human γ-globulins as follows: 100×[1-(number of plaques after incubation with human γ-globulins)/(number of plaques after incubation without human γ-globulins)]. Each value is the mean ± standard error of the results of 6 independent experiments. The statistical analysis was performed by one-way ANOVA followed by the Dunnett multiple comparisons test. n.s., not significant.

**Supplementary Fig. 7.**
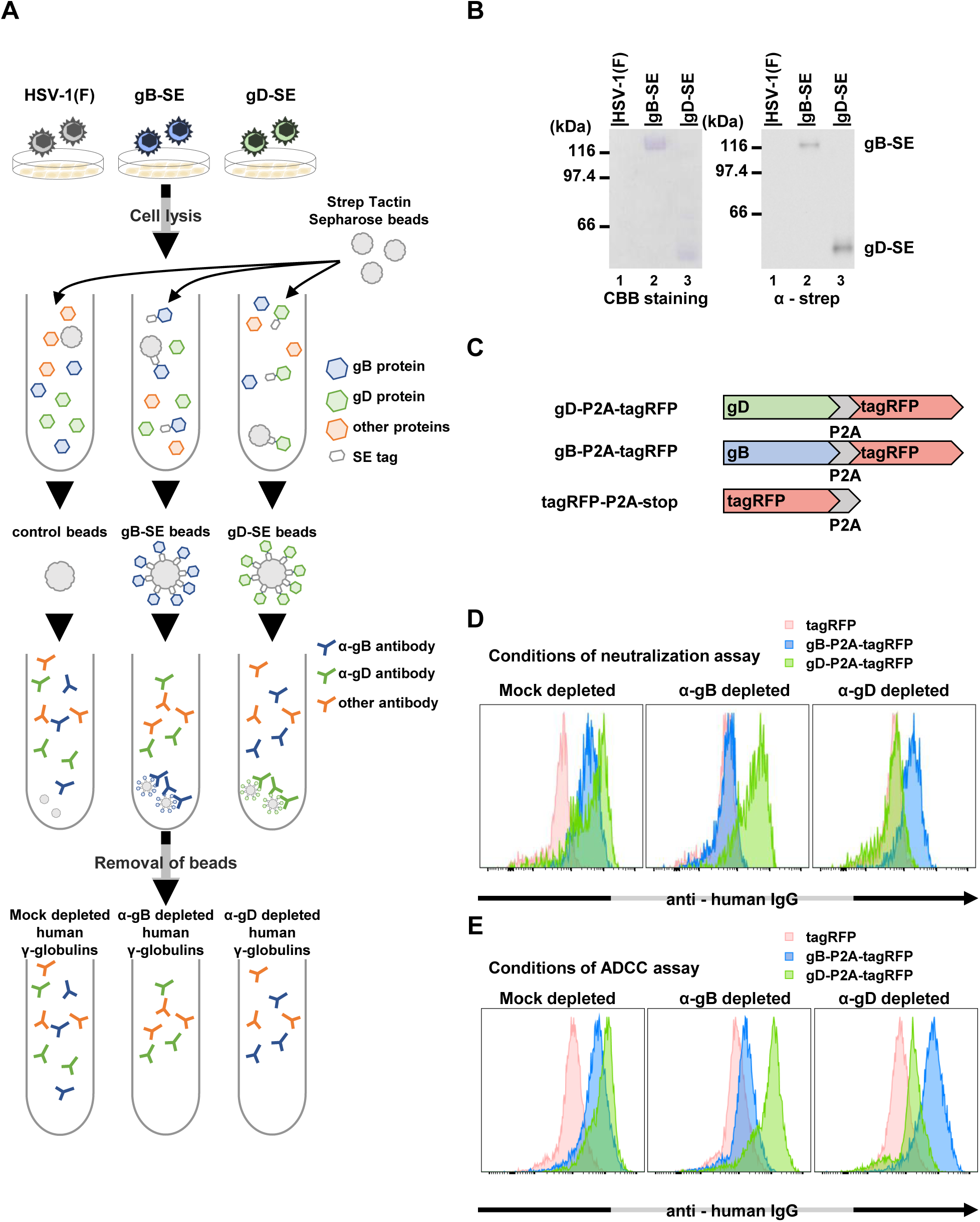
Depletion of gB or gD antibodies from pooled human γ-globulins. (A) Schematic diagram of depletion of gB or gD antibodies from human γ-globulins. Vero cells were infected with wild-type HSV-1(F), YK717 (gB-SE), or YK718 (gD-SE) at an MOI of 0.5 for 24 h. The cells were then lysed and incubated with Strep-Tactin Sepharose beads. For mock-depletion, human γ-globulins (0.082 and 3.04 mg/mL for neutralizing and ADCC assays, respectively) were incubated with Strep-Tactin Sepharose beads previously incubated with the lysate of HSV-1(F)-infected cells. For depletion of gB or gD antibodies from human γ-globulins, human γ-globulins were incubated with gB-SE or gD-SE immobilized on Strep-Tactin Sepharose beads. (B) The Strep-Tactin Sepharose beads reacted with lysates of infected cells as described in A were divided into 2 aliquots. One aliquot was analyzed by electrophoresis in a denaturing gel and stained with Coomassie brilliant blue (CBB) (left gel), and the other aliquot was analyzed by immunoblotting with anti-strep antibody (right gel). (C to E) HEK293FT cells were transfected with pcDNA3.1-tagRFP-P2A-stop, pcDNA3.1-gB-P2A-tagRFP, or pcDNA3.1-gD-P2A-tagRFP, encoding tagRFP-P2A-stop, gB-P2A-tagRFP, or gD-P2A-tagRFP, respectively (C). At 48 h post-transfection, cells were incubated with mock-depleted human γ-globulins (mock-depleted) or human γ-globulins depleted with anti-gB-SE (anti-gB depleted) or anti-gD-SE (anti-gD depleted), and tagRFP+ cells were analyzed (D and E). The data are representative of 3 independent experiments (B and D). anti, α.

**Supplementary Fig. 8.**
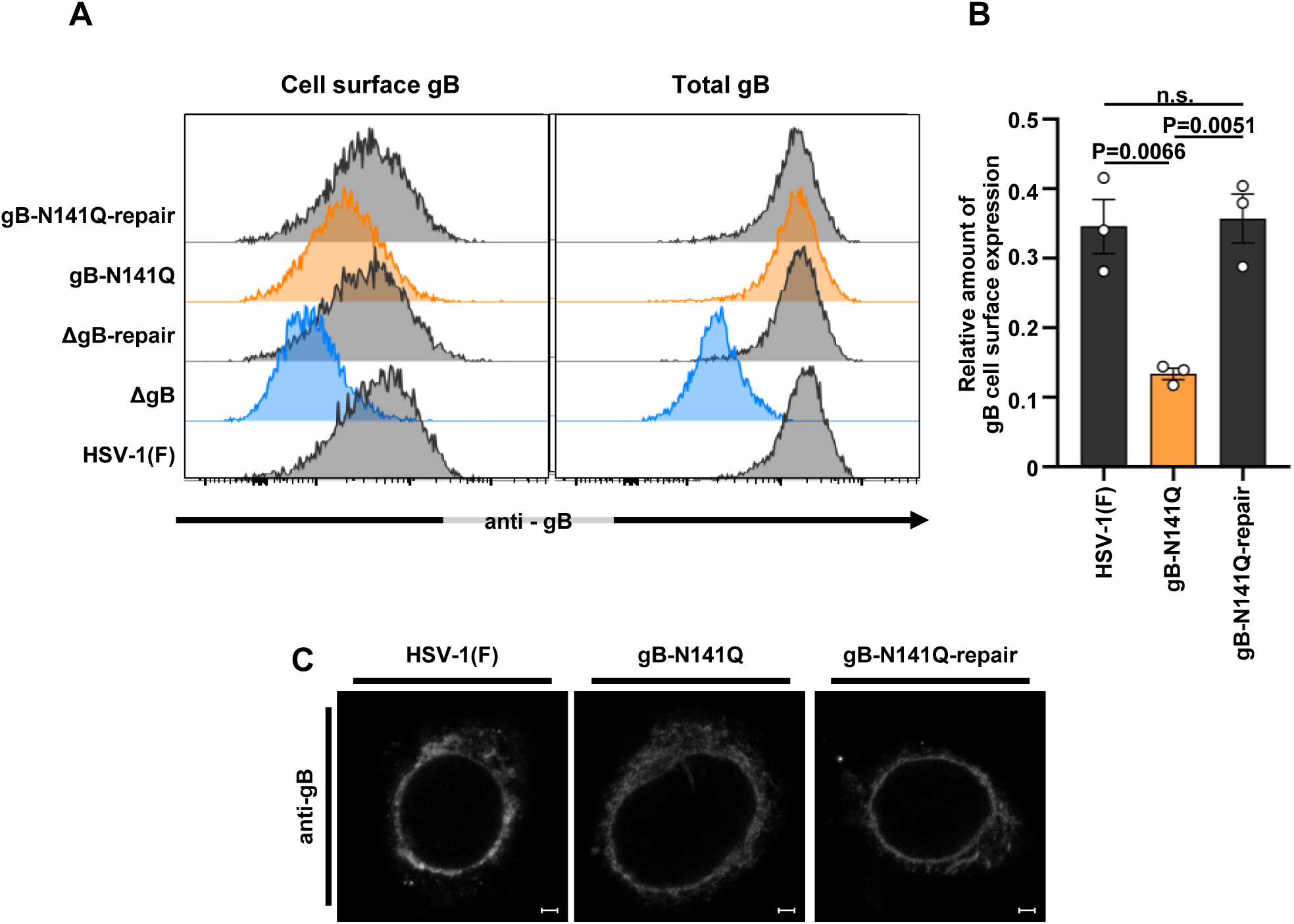
Effect of N-glycan at gB Asn-141 on cell surface expression of gB in HSV-1-infected cells. (A) Vero cells were infected with wild-type HSV-1(F), gB-N141Q, gB-N141Q-repair, ΔgB, or ΔgB-repair at an MOI of 1. At 24 h post-infection, cell surface expression (left panel) and total expression (right panel) of gB in infected cells were analyzed by flow cytometry. (B) Quantitative bar graph of the cell surface expression of gB shown in (A). The relative amount of expression of gB on the cell surface was calculated as follows: [(mean fluorescent intensity for gB expression on the surfaces of cells infected with the indicated virus) − (mean fluorescent intensity for gB expression on the surfaces of cells infected with ΔgB)]/[(mean fluorescent intensity for total gB expression in cells infected with the indicated virus) − (mean fluorescent intensity for total gB expression in cells infected with ΔgB)]. (C) Vero cells infected with wild-type HSV-1(F), gB-N141Q, or gB-N141Q-repair at an MOI of 5 for 18 h were fixed, permeabilized, stained with antibody to gB, and examined by confocal microscopy. Scale bars = 2 μm. The data are representative of 3 independent experiments (A and C). Each value is the mean ± standard error of the results of 3 independent experiments (B). Statistical analysis was performed by one-way ANOVA followed by the Tukey test. n.s., not significant (B).

**Supplementary Fig. 9.**
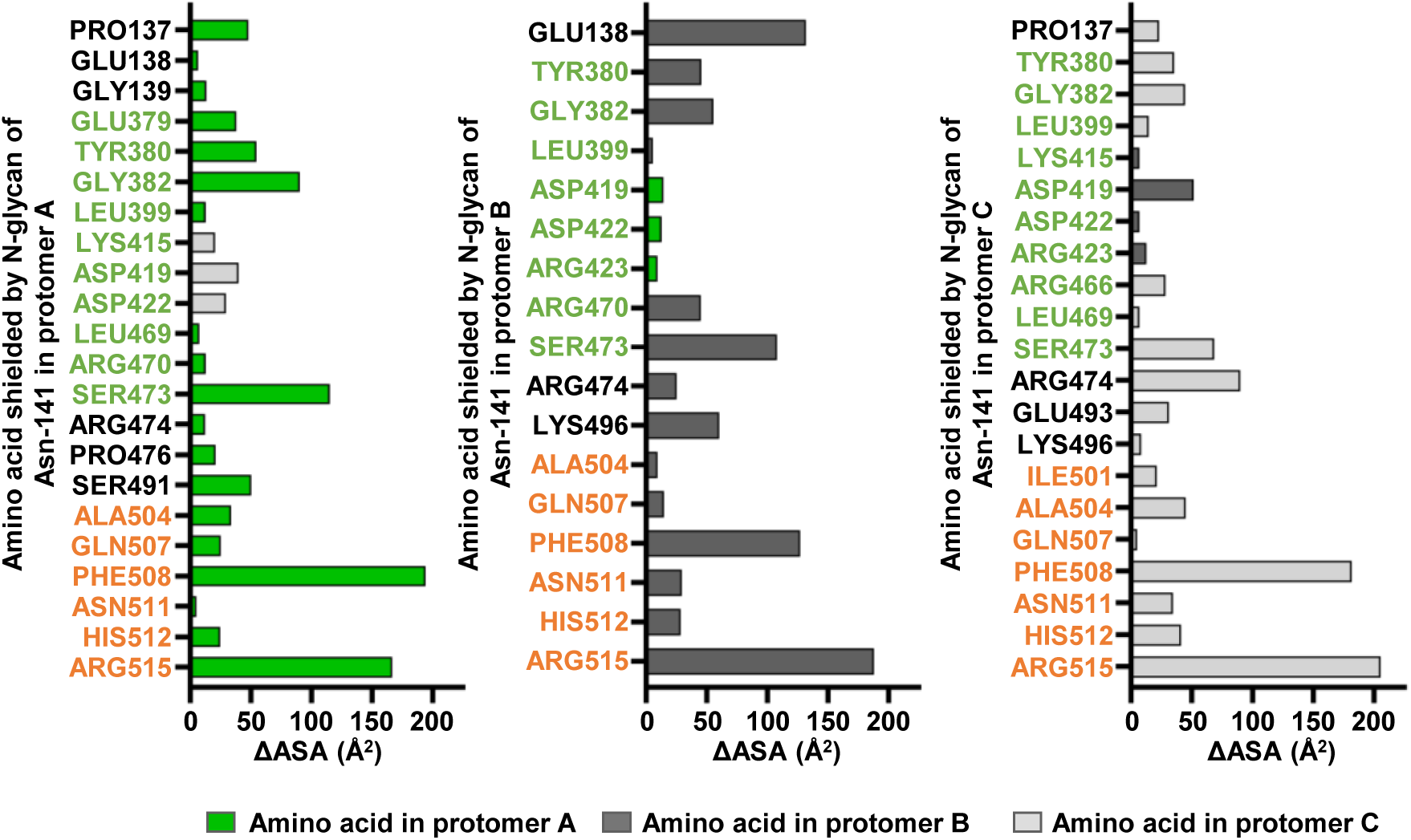
Bar graphs showing effects of N-glycan shield at gB Asn-141 on the antigenicity of gB in the prefusion state. Bar graphs showing ΔASA values (> 5 Å^2^) for each of the protomers of the gB trimer in the prefusion state. Abbreviations of the amino acids residues mapped to FR2 and FR3 (20, 21) are colored in green and orange, respectively.

**Supplementary Table 1A.**
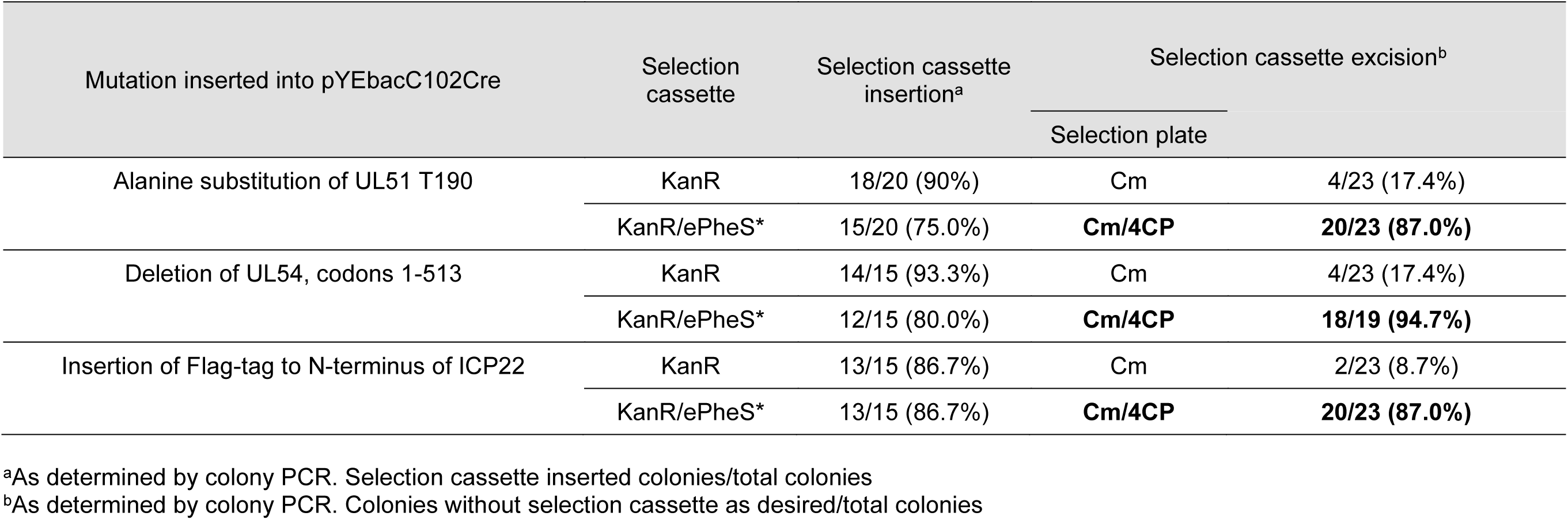
Efficiency of two-step Red-mediated recombination with ePheS* cassette.

**Supplementary Table 1B.**
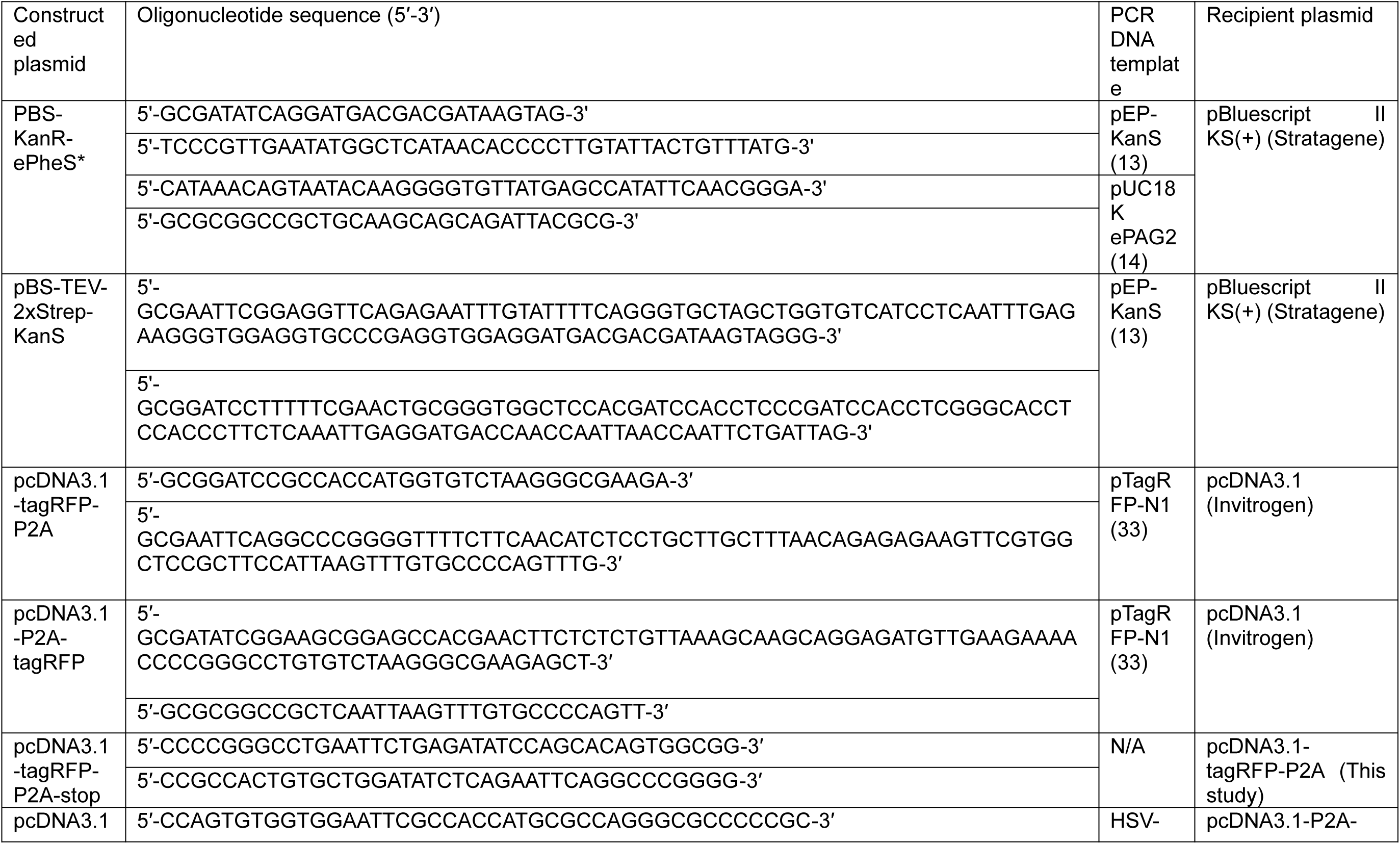
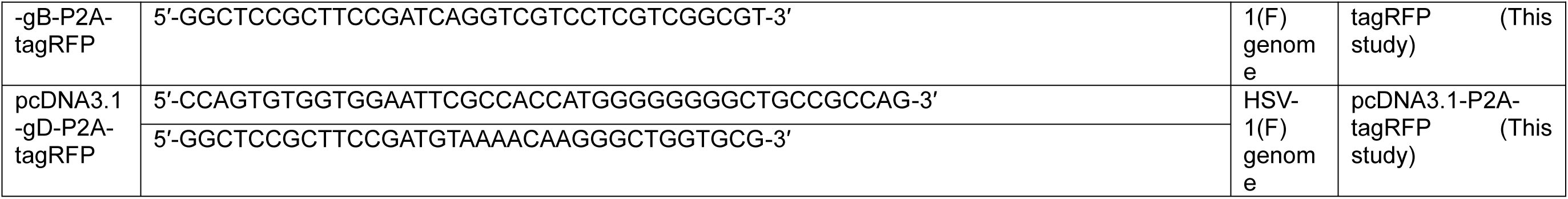
Oligonucleotide sequences and DNA templates for the construction of plasmids.

**Supplementary Table 1C.**
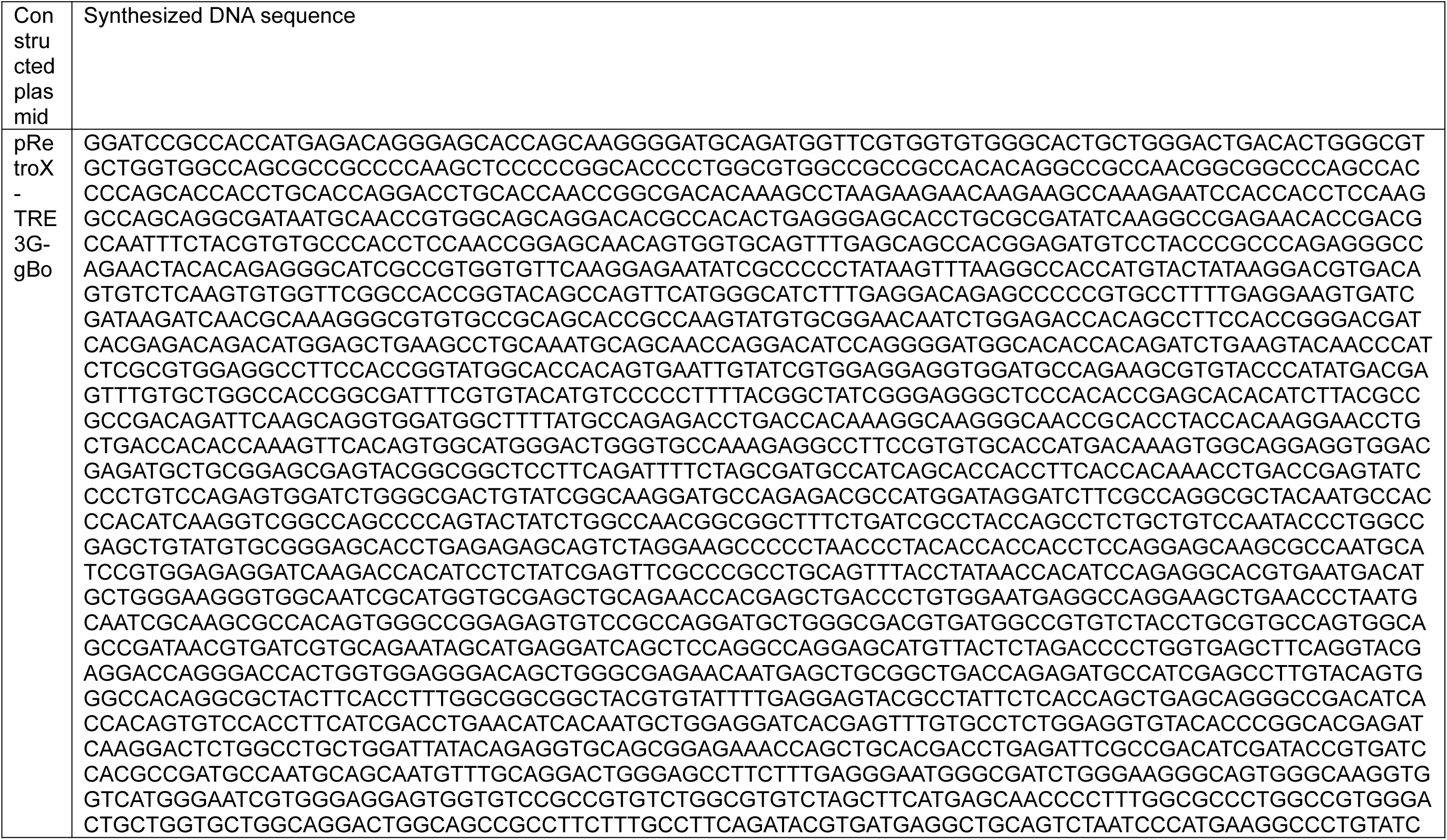
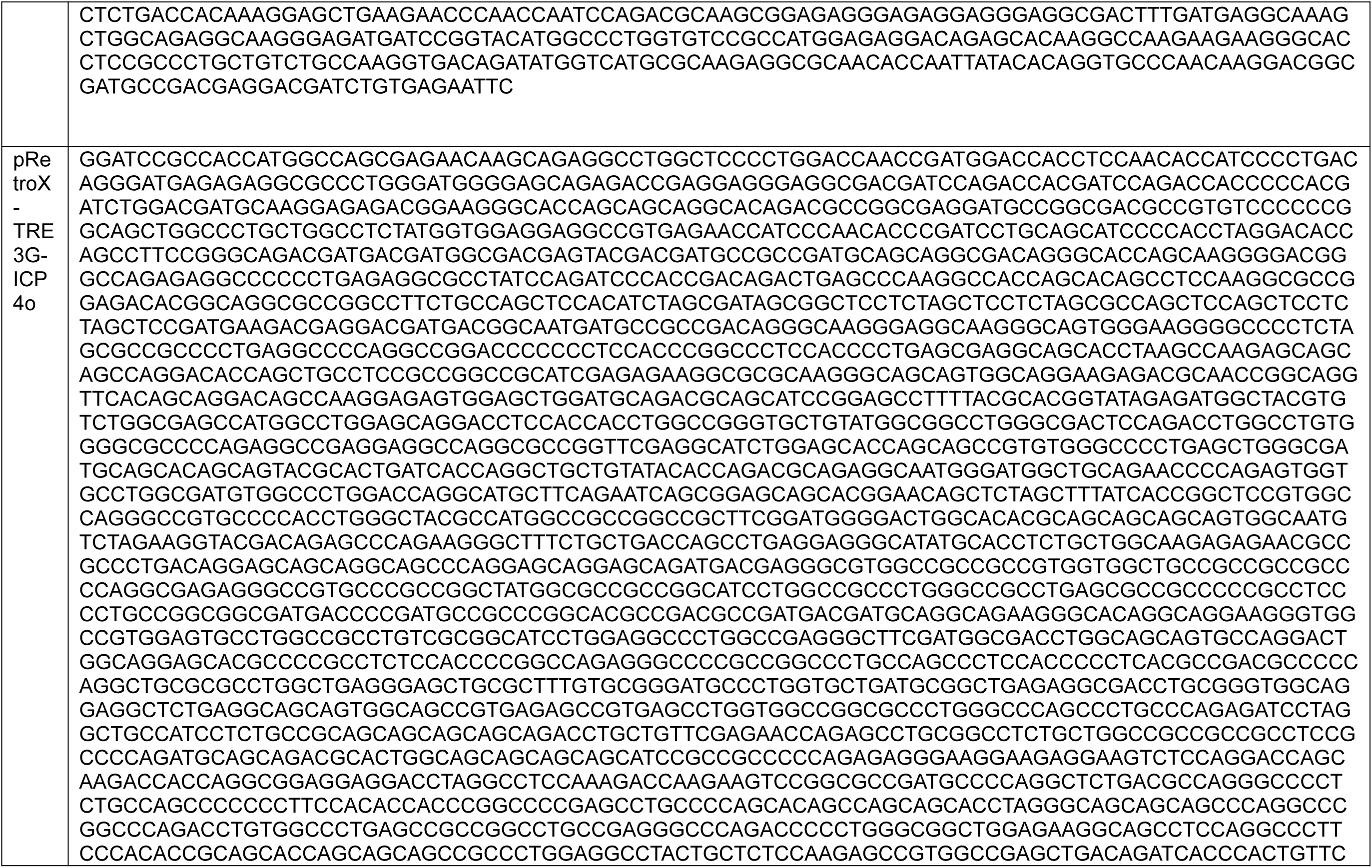
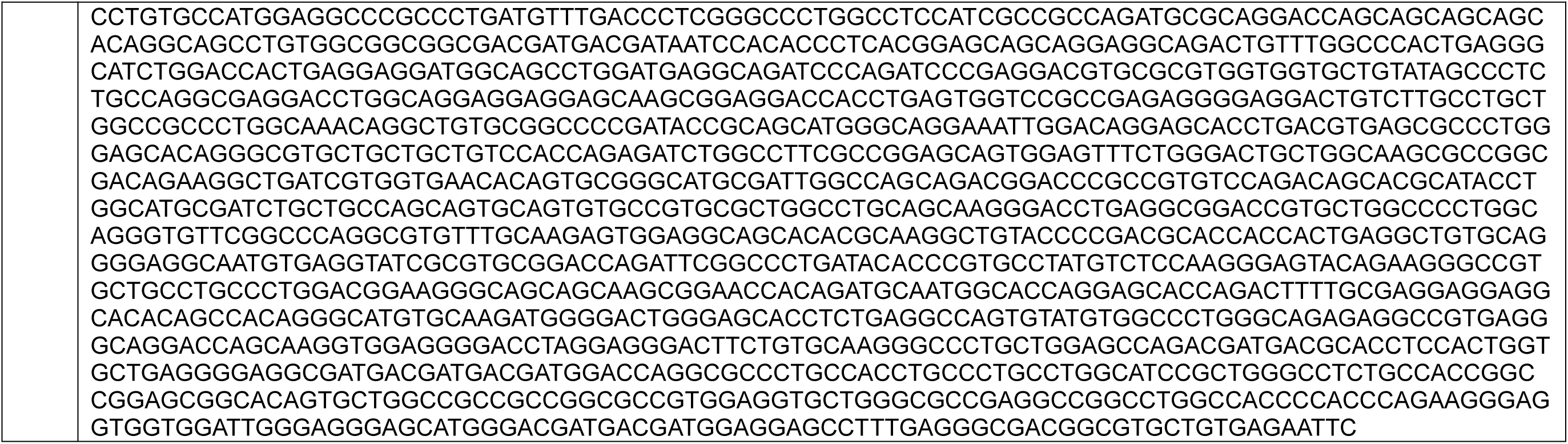
Summary of synthesized plasmids.

**Supplementary Table 1D.**
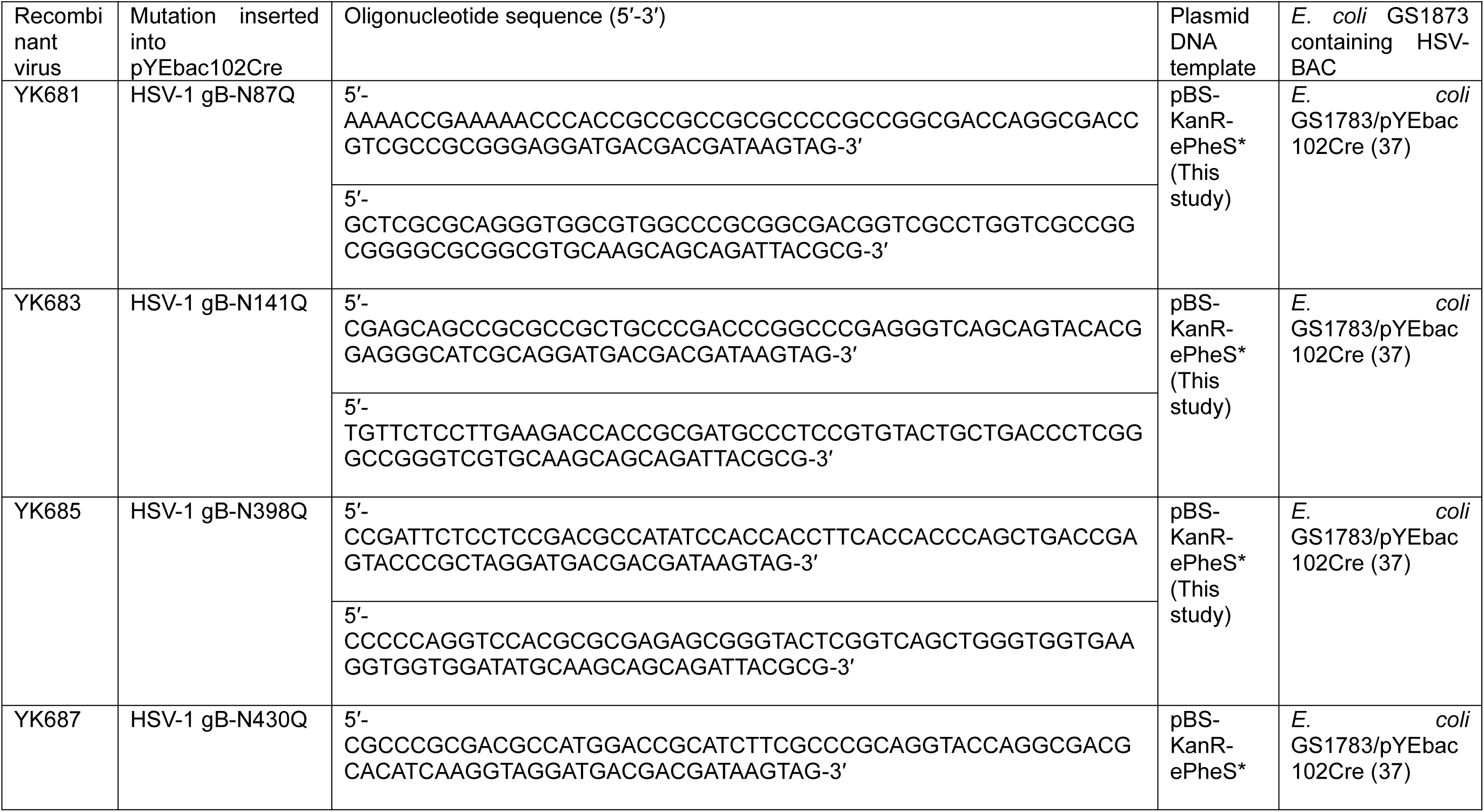
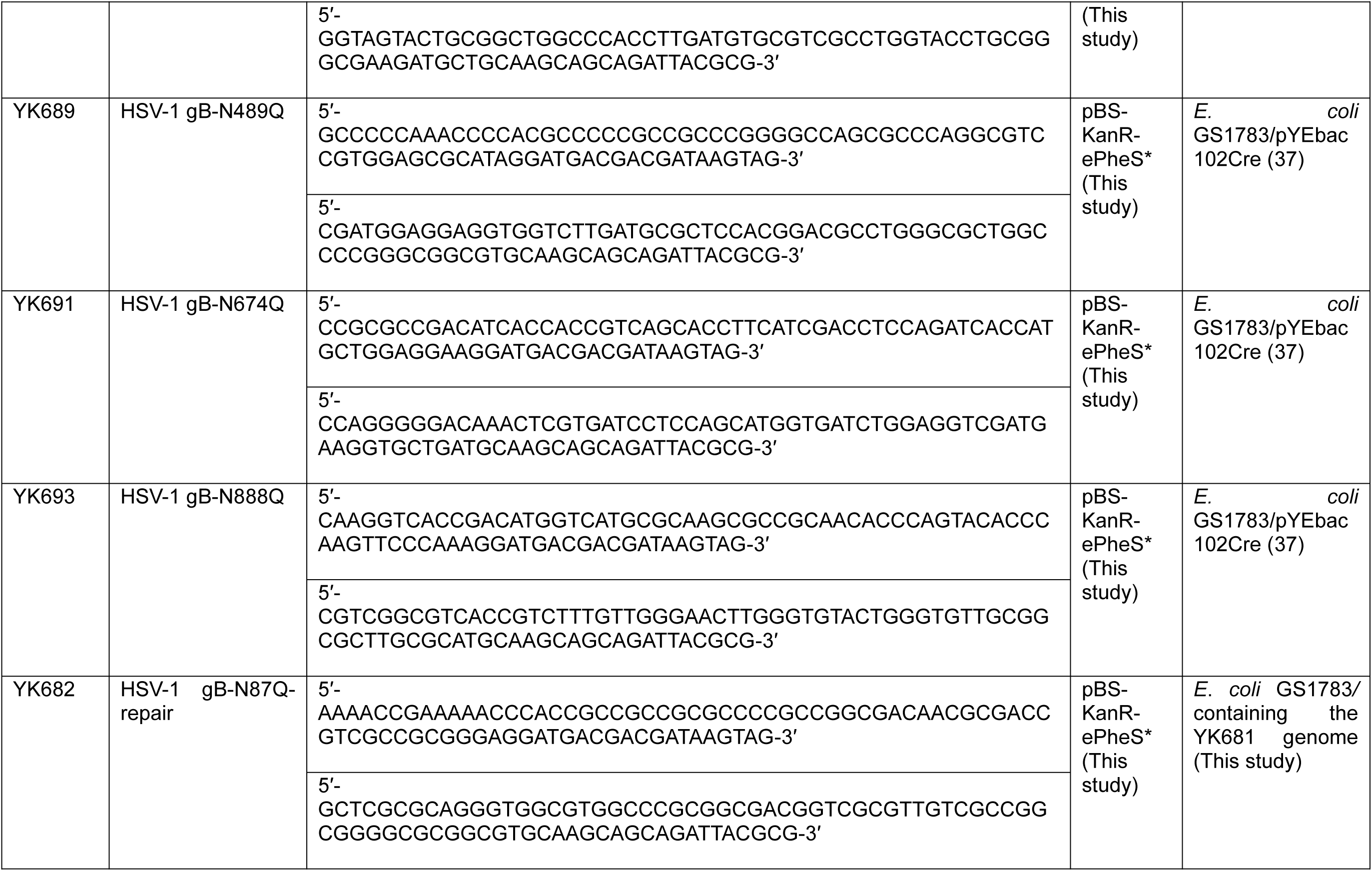
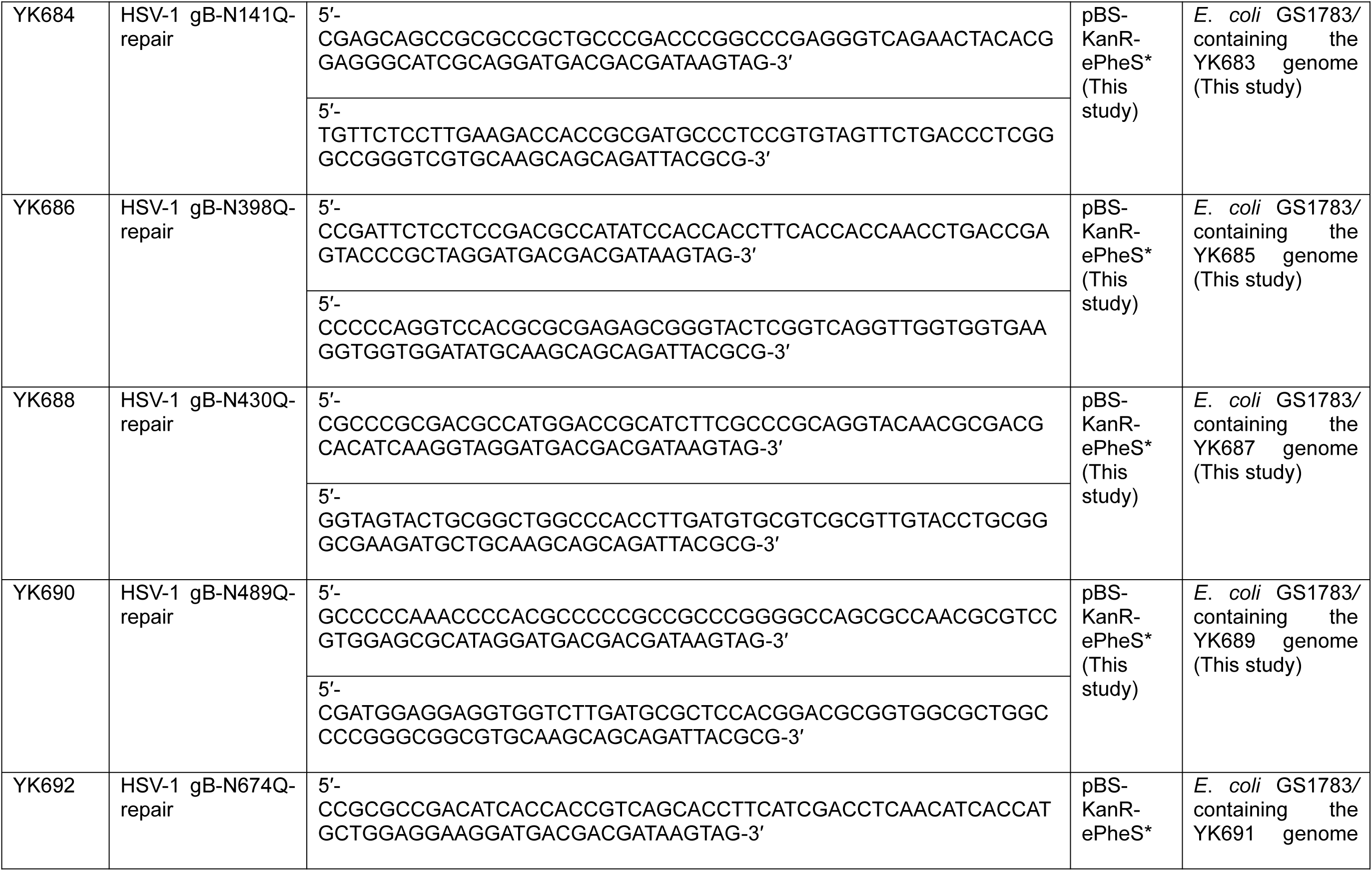
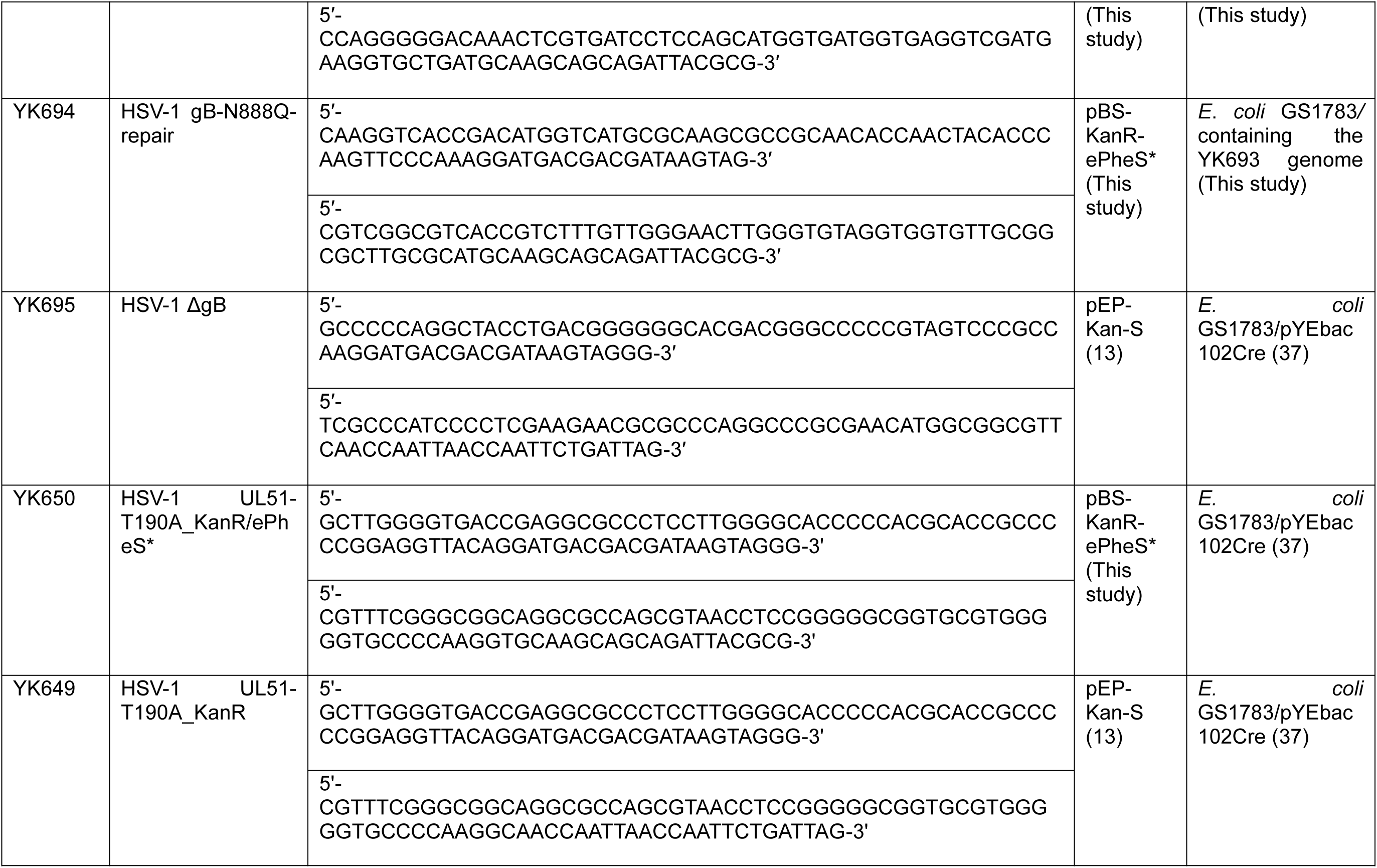
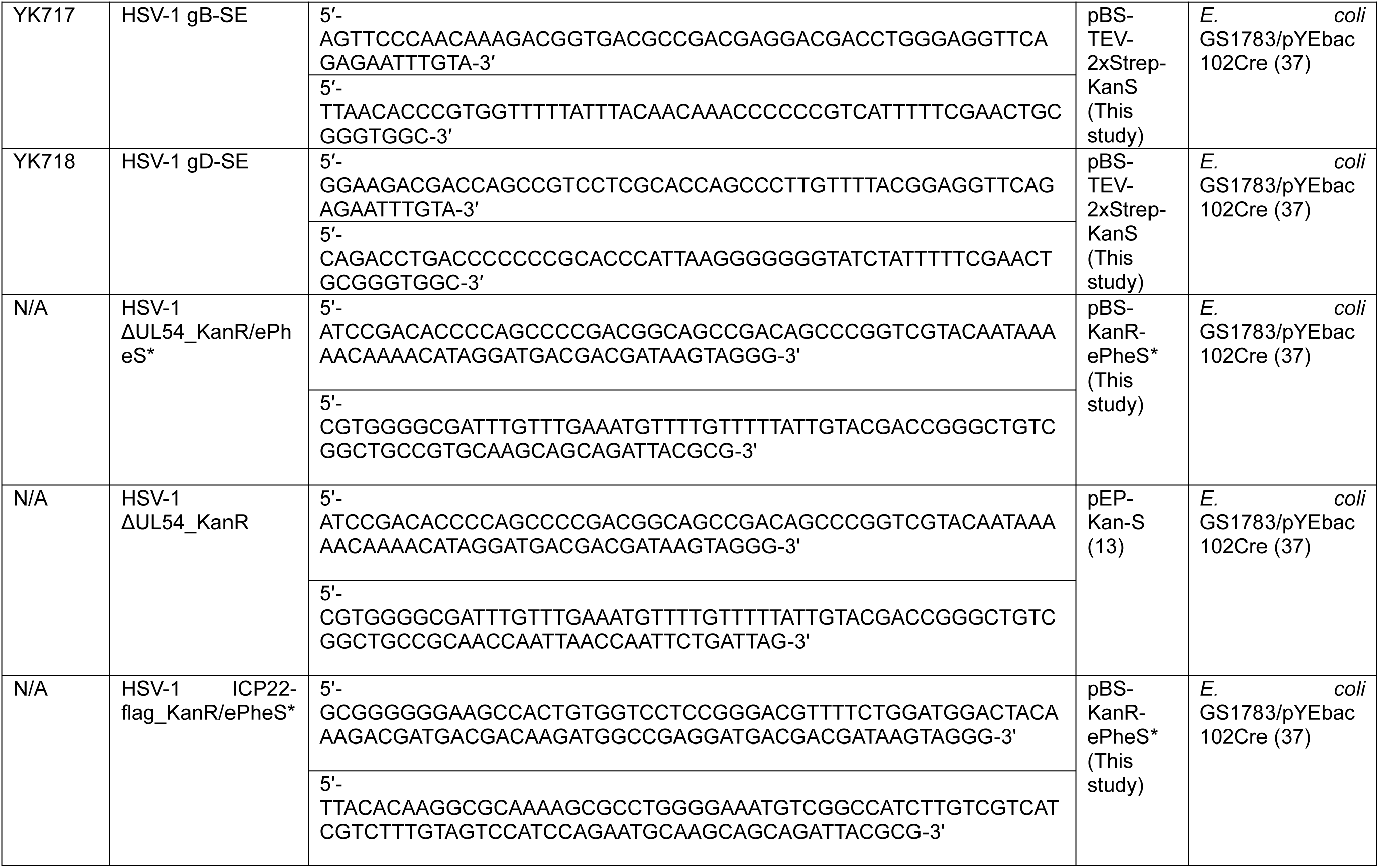
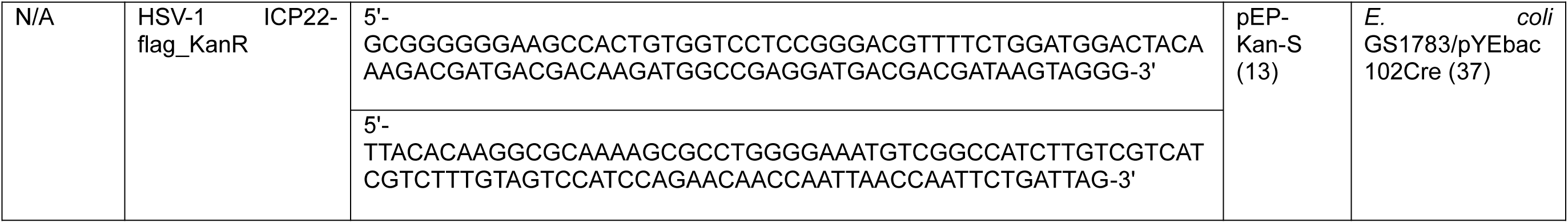
Oligonucleotide sequences, plasmid, and E. coli GS1873 containing HSV-BAC for the construction of recombinant viruses or mutagenesis.

